# African rice (*Oryza glaberrima*) genomic introgressions impacting upon panicle architecture in Asian rice (*O. sativa*) lead to the identification of key QTLs

**DOI:** 10.1101/2023.04.25.538245

**Authors:** Hélène Adam, Andrés Gutierrez, Marie Couderc, François Sabot, Fabrice Ntakirutimana, Julien Serret, Julie Orjuela, James Tregear, Stefan Jouannic, Mathias Lorieux

## Abstract

**Background:** Developing high yielding varieties is a major challenge for breeders tackling the challenges of climate change in agriculture. The panicle (inflorescence) architecture of rice is one of the key components of yield potential and displays high inter- and intra-specific variability. The genus Oryza features two different crop species: Asian rice (*Oryza sativa* L.) and the African rice (*O. glaberrima* Steud). One of the main morphological differences between the two independently domesticated species is the structure (or complexity) of the panicle, with *O. sativa* displaying a highly branched panicle, which in turn produces a larger number of grains than that of *O. glaberrima*. The genetic interactions that govern the diversity of panicle complexity within and between the two species are still poorly understood.

**Results:** To identify genetic factors linked to panicle architecture diversity in the two species, we used a set of 60 Chromosome Segment Substitution Lines (CSSLs) issued from third generation backcross (BC_3_DH) and carrying genomic segments from *O. glaberrima* cv. MG12 in the genetic background of *O. sativa* Tropical Japonica cv. Caiapó. Phenotypic data were collected for rachis and primary branch length, primary, secondary and tertiary branch number and spikelet number. A total of 15 QTLs were localized on chromosomes 1, 2, 3, 7, 11 and 12 and QTLs associated with enhanced secondary and tertiary branch numbers were detected in two CSSLs. Furthermore, BC_4_F_3:5_ lines carrying different combinations of substituted segments were produced to decipher the effects of the identified QTL regions on variations in panicle architecture. A detailed analysis of phenotypes versus genotypes was carried out between the two parental genomes within these regions in order to understand how *O. glaberrima* introgression events may lead to alterations in panicle traits.

**Conclusion:** Our analysis led to the detection of genomic variations between *O. sativa* cv. Caiapó and *O. glaberrima* cv. MG12 in regions associated with enhanced panicle traits in specific CSSLs. These regions contain a number of key genes that regulate panicle development in *O. sativa* and their interspecific genomic variations may explain the phenotypic effects observed.

## Background

Improving or enhancing the sustainability of rice yields continues to be crucial challenge in the breeding of this crop, especially in the context of a growing world population and with regard to climate change. Inflorescence architecture in rice directly affects yield potential through the regulation of grain number and harvesting ability. Grain number per panicle depends on orders and numbers of branches (i.e., primary, secondary and potentially tertiary branches as well as both lateral and terminal spikelets) and on the length of each axis. Rice panicle architecture is based on the production of a series of lateral meristems with distinct identities (Itoh et al., 2005). After the floral transition, the Shoot Apical Meristem (SAM) is converted into an indeterminate Rachis Meristem (RM) which produces several axillary meristems, the primary branch meristems (PBMs) that subsequently give rise to primary branches (PBs). Once the RM becomes degenerate (i.e., it has lost its activity), the newly formed PBM elongates and initiates a variable number of axillary meristems forming a new branch order. These acquire the identity of secondary branch meristems (SBMs) and develop into secondary branches (SBs), which in turn may produce tertiary branch meristems (TBMs), or alternatively differentiate directly into lateral spikelet meristem (SpMs) and then florets. The basic architecture of the panicle is thus determined by patterns of axillary meristem formation and the specification of their identities. Overall, the complexity and diversity of panicle branching can be considered as governed by two key elements: the number of axillary meristems produced during the indeterminate phase; and the rate of meristem fate transition, which determines whether an axillary meristem grows into a higher-order branch or differentiates into a spikelet.

Over recent decades, a number of genes and quantitative trait loci (QTLs) associated with panicle development and affecting its architecture have been identified and functionally described in rice (Xing and Zhang 2010; Chongloi et al., 2019; Yin et al., 2021; Wang et al., 2021). Typically, these genes and QTLs are implicated in axillary meristem formation or in the conversion of indeterminate meristems to spikelets, thus influencing the branching complexity of the panicle (Komatsu et al., 2003a; Komatsu et al., 2003b; Kobayashi et al., 2009; Yoshida et al., 2013; Bai et al., 2016a).

Most studies of rice panicle development have focused on the Asian crop plant *Oryza sativa* L. However, a second species of cultivated rice, *O. glaberrima* Steud, was domesticated independently of Asian rice, in the inner delta of Niger river (Cubry et al., 2018) from the wild relative species *O. barthii* around 2,500 years ago. Cultivation of *O. sativa* was introduced into West Africa around 400 years ago and this crop has since largely replaced *O. glaberrima* – although the latter is still grown, it represents only 1-2% of the total cultivated area. *O. glaberrima* is more adapted to a variety of ecologically and climatically diverse regions, its agronomically useful characters including higher tolerance to drought, high temperatures and infertile soils as well as a greater resistance to various biotic stresses (Jones et al., 1997; Linares, 2002; Thiémélé et al., 2010; Wambugu et al., 2013; Li et al., 2015; Wambugu et al., 2021; Lorieux et al., 2013). While *O. sativa* is less adapted to the African environment, this species has a higher yield potential than *O. glaberrima*, the difference being partly explained by the higher panicle complexity of Asian rice (Harrop et al., 2019). Even if molecular pathways associated with the regulation of axillary meristem identity seem to be conserved between the two species, it was possible to identify a set of genes displaying differential expression in relation to panicle phenotypic variability between African and Asian rice (Harrop et al., 2019). Moreover, a Genome Wide Association Study (GWAS) carried out on a panel of African rice accessions allowed the identification of several new genomic regions associated with panicle branching diversity and climatic variables (Cubry et al., 2020).

In the context of climate change, achieving high tolerance to environmental stresses is of paramount importance to rice agriculture while knowledge relating to the genetic control of traits governing yield potential in African rice remains very limited. Agronomic traits such as heading date, yield, plant height and grain size are controlled by many genes exerting major and/or minor effects. Identifying the genes and genomic regions associated with interspecific variation in the number of grains produced per panicle is challenging due to the polygenic nature and environmental sensitivity of panicle branching traits.

Libraries of introgression lines (ILs) provide a useful means to investigate genetic phenomena from an interspecific perspective. An introgression line library is a collection of lines with a common genetic background (the recipient genome), each carrying one or a few genomic regions originating from a donor genome. The ILs are generally obtained through recurrent backcrossing onto the recipient parent, followed by several rounds of self-fertilization (BC_n_F_n_) or double haploidization (BC_n_DH). The genome of the donor genotype is then represented in the library by discrete homozygous chromosomal fragments. Such ILs are also called Chromosome Segment Substitution Lines (CSSLs). In contrast to segregating populations, such as Recombinant Inbred Lines (RILs), Doubled Haploid (DH), Backcross (BC) or F_2_:_3_ populations, QTL analyses using CSSLs are suitable to evaluate minor allelic differences conferred by additive QTLs in a uniform genetic background in a reduced population size (Eshed and Zamir, 1995; Koumproglou et al., 2002). Genetic dissection of complex traits can thus be achieved by combining genetic variation with introgressed genomic fragments so as to reduce interference effects between QTLs. Hence, CSSLs provide a powerful means to identify QTLs for complex traits with minor and/or additive effects (Balakrishnan et al., 2019; Zhang et al., 2022). This approach may provide access to novel and potentially beneficial genes “hidden” in the genetic background of a related species that can be discovered when placed in the genetic background of a cultivated species.

In the present study, we evaluated several panicle morphological traits in a BC_3_DH CSSL population developed from an interspecific cross between the recipient parent *O. sativa ssp. japonica* (cv. Caiapó) and the donor parent *O. glaberrima* (cv. MG12) (Table 1) (Gutiérrez et al., 2010). QTLs relating to modifications in panicle morphology were mapped on the rice genome, and 4^th^ generation backcross (BC_4_F_3:5_) generations were produced to evaluate the effect of each region on panicle phenotype variation. Finally, DNA polymorphisms between the two parental genomes were investigated in detail for some key genes that were previous implicated in panicle branching diversity in *O. sativa*. The results obtained will help allow the identification of functionally significant polymorphisms and, more generally, provide an insight into the genetic bases of panicle architecture diversity between the two species.

**Table 1:** Panicle Traits analyzed in the CSSLs

| Variable | Name | Measurement unit |
| --- | --- | --- |
| PBN | Primary branch number | Number/panicle |
| SBN | Secondary branch number | Number/panicle |
| SBN/PBN | Secondary branch number per Primary branch num-<br>ber | Number/panicle |
| TBN | Tertiary branch number | Number/panicle |
| SpN | Spikelet number | Number/panicle |
| PBL | Primary Branch length | cm/panicle |
| RL | Rachis Length | cm/panicle |

## Results

### Panicle Traits in the CSSL population

In order to identify new genetic factors governing the panicle branching diversity observed between *O. sativa* cv. Caiapó (referred to as *Os_Caiapó* hereafter) and *O. glaberrima* cv. MG12 (referred to as *Og_MG12* hereafter), phenotypic measurements were performed on a population of 60 BC_3_DH CSSLs and their parents (Additional file1: Table S1) (Gutiérrez et al. 2010).

Six quantitative traits were evaluated per panicle: rachis length (RL); primary, secondary and tertiary branch numbers (PBN, SBN, TBN respectively); primary branch length average (PBL); and spikelet number (SpN) (Table1, Additional file1: Table S2). The two parents showed contrasting panicle phenotypes with a higher panicle complexity observed in *Os_Caiapó* compared to *Og_MG12*. More specifically, the *Os_Caiapó* parent displayed panicles with more PBN and SBN leading to a higher SpN compared to *Og_MG12* (Fig. 1A). In contrast, the *Og_MG12* parent produced a panicle with longer PBs compared to *Os_Caiapó*. Since the trials were conducted as two repeated experiments, we computed broad sense heritability for each experiment (Additional file1: Table S3). All the measured variables showed broad sense heritability values higher than 0.8 except for TBN. For all traits, the mean of the CSSL population was similar to the mean of the recurrent *Os_Caiapó* parent (Additional file1: Table S4). The coefficient of variation for the SBN/PBN ratio was higher in the CSSL population than in *Os_Caiapó* (26.10% versus 18.26%). Histograms of the CSSLs for panicle traits showed a continuous distribution for all traits except TBN (Fig. 1B). The observed distributions were similar between the two repeated experiments. The abnormal distribution of the TBN trait was associated with a high value of CV (Coefficient Variation) and a low heritability value. This observation probably results from the rare and unstable nature of this trait which is dependent on environmental conditions and does not appear frequently in the *Os_Caiapó* parent. Moreover, the formation of tertiary branches has not been described in *O*. *glaberrima* populations (Cubry et al., 2020), nor was it observed in our earlier experiments.

**Fig. 1.**
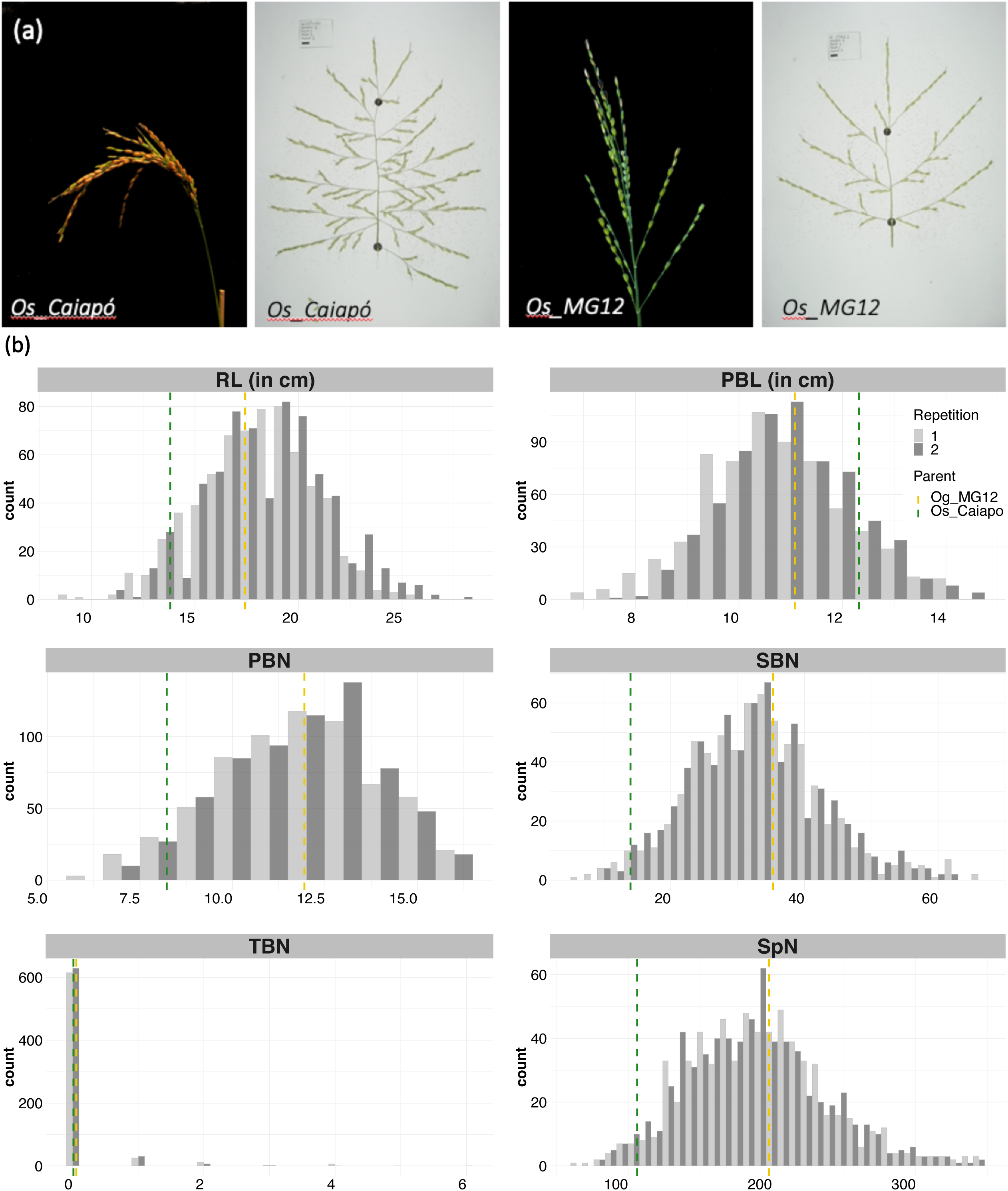
Panicle structure and morphological panicle trait values in the two parental lines and the 60 BC3DH CSSLs. **(a)** Panicle Architecture contrasts between the two parental lines (*Os_Caiapó* and *Og_MG12*). **(b)** Distribution of panicle trait values in the 60 BC3DH CSSLs. Dashed vertical lines correspond to the mean values of each parent (green for *Og_MG12* and yellow for *Os_Caiapó*). Values for repetitions 1 and 2 are shown in light gray and dark gray, respectively. Abbreviations: RL, Rachis Length; PBN, Primary Branch Number; PBL, Primary Branch Length; SBN, Secondary Branch Number; TBN, Tertiary Branch Number; SpN, Spikelet Number.

By analyzing relationships between the different panicle traits, we observed a strong correlation (0.95) between SpN and SBN (Fig. 2A). Other traits were moderately correlated with each other, notably SBN and PBN. We also observed a low correlation between PBL and SBN. Principal component analysis showed a separate distribution of *Os_Caiapó* and *Og_MG12* parents on the PC1 and PC2 axes. In contrast, we observed an overlapping of the BC_3_DH lines with the *Os_Caiapó* recurrent parent (Fig. 2B). Analysis of variable contributions showed that SpN is the main trait contributing to the diversity observed in this population.

**Fig. 2.**
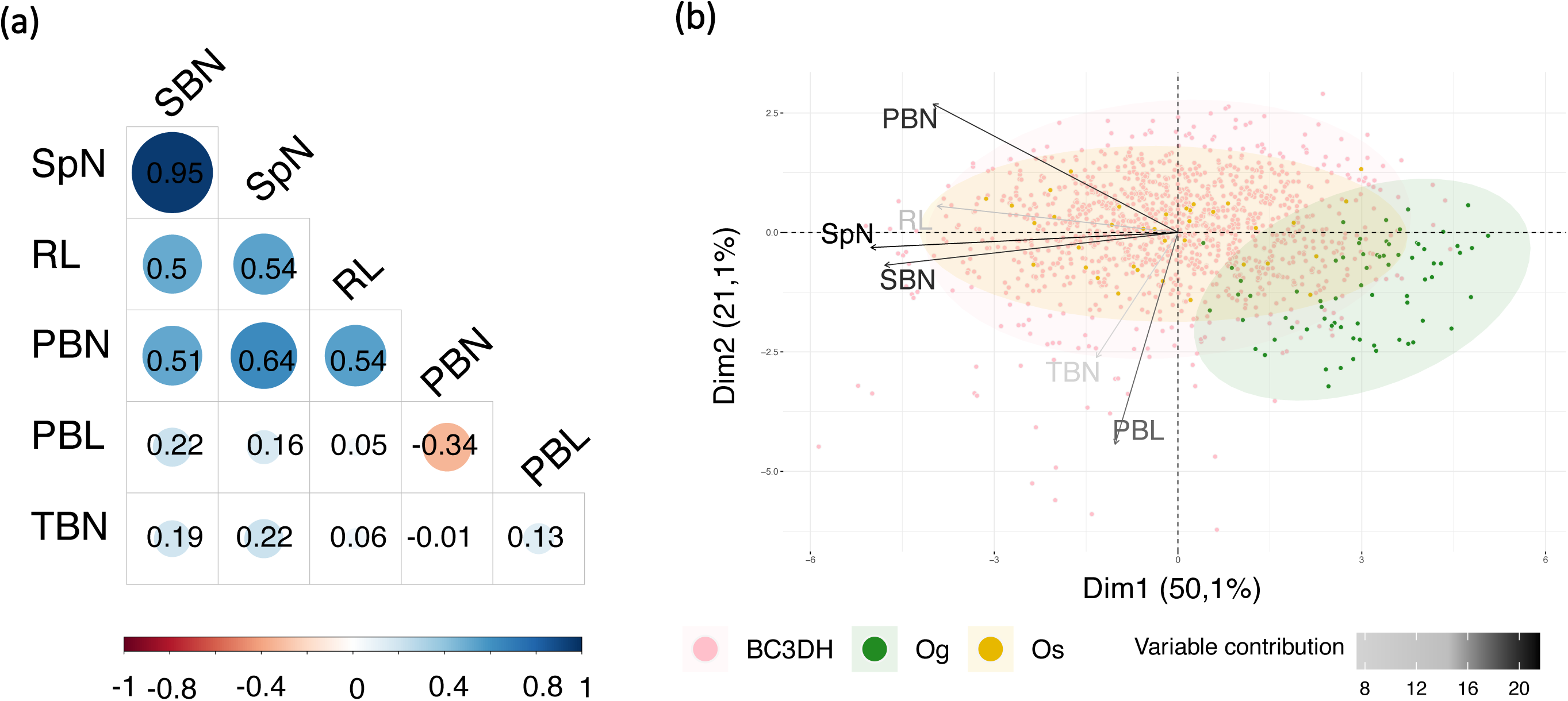
Description of the morphological panicle traits observed in the BC3DH CSSL lines and parents. **(a)** Correlations between panicle traits. **(b)** Principal component analysis with traits contributions and individual panicle distributions. Abbreviations: RL, Rachis Length; PBN, Primary Branch Number; PBL, Primary Branch Length; SBN, Secondary Branch Number; TBN, Tertiary Branch Number; SpN, Spikelet Number.

### Detection of QTLs associated with panicle architecture traits

The Dunnett’s test revealed a total of 37 lines with significant differences compared to the *Os_Caiapó* recurrent parent for all traits combined (Additional file1: Table S5). In general, these lines bear at least two *Og_MG12* introgression fragments. For this reason, assigning the underlying QTL to a single chromosome segment was not straightforward. The CSSL Finder software was therefore used to detect genomic regions associated with panicle phenotype variation with a F-test value at each marker (Gutiérrez et al., 2010). Graphical genotype representations were also produced in which the CSSLs were ordered by trait value for each evaluated trait (Additional file 2: Figure S1). We considered the F-test as significant when its value was higher than 10.0. QTLs were assigned to the *Og_MG12* introgressed regions of these CSSLs if the F-test was significant and confirmed by the graphical genotype logical analysis.

In total, 15 QTLs were detected for all combined traits analyzed with the exception of rachis length (Fig. 3). The F-test value of the markers allowing the detection of each QTL are reported in Table 2 along with their positions in the *O. sativa* and *O. glaberrima* reference genomes (cv. Nipponbare IRGSP-1.0 and cv. CG14 OglaRS2 respectively).

**Fig. 3.**
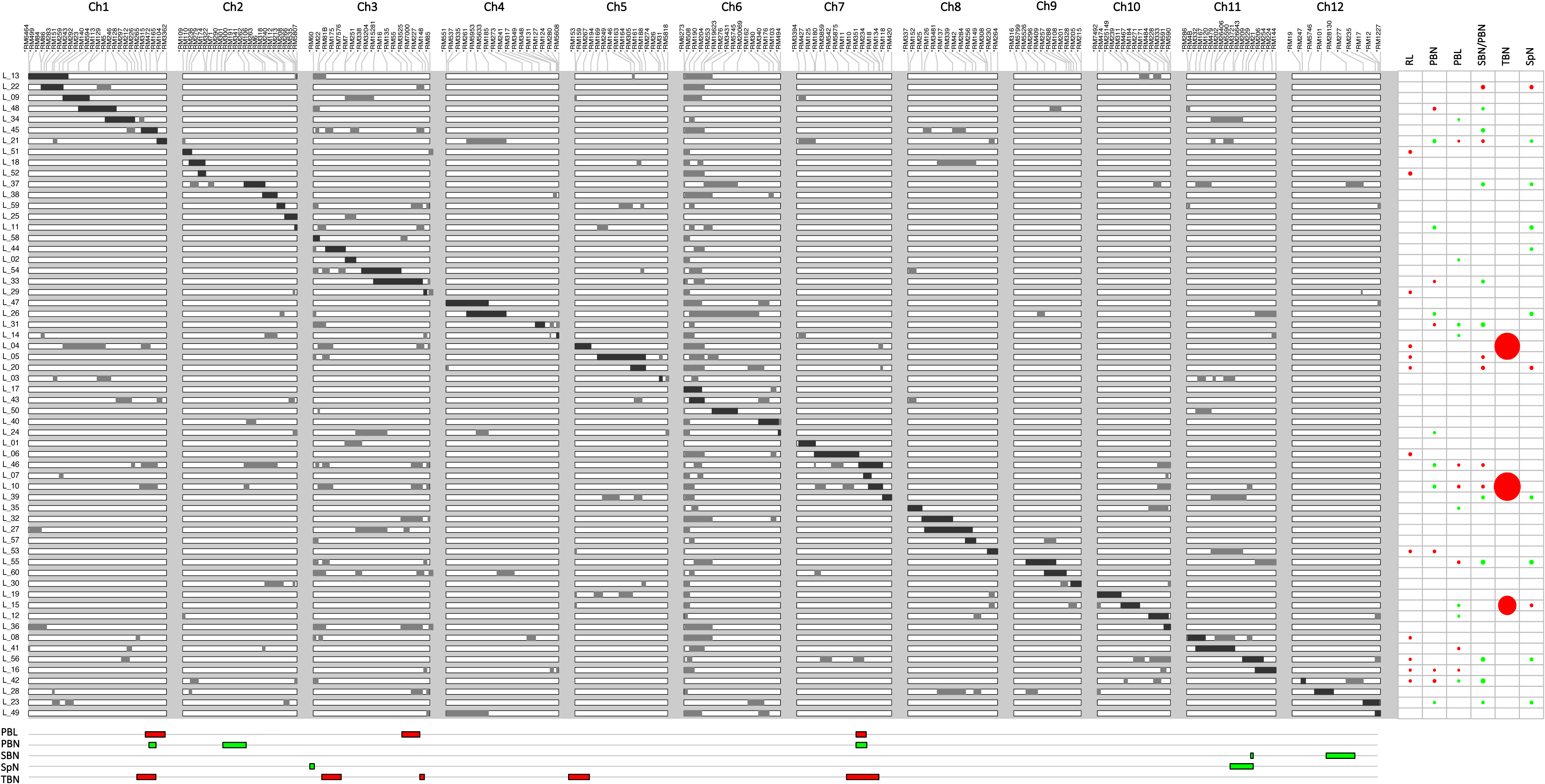
Graphic representation of the genotypes of the 60 BC3DH CSSLs, showing line x trait significant associations and QTL positions. The 12 chromosomes are covered by 200 evenly dispersed SSR markers. Genotypes are displayed horizontally. Black areas represent the *O. glaberrima* MG12 target chromosome segments, the set of segments broadly covering the entire *O. glaberrima* MG12 genome. White areas represent the *O. sativa Caiapó* genetic background and grey areas represent *O. glaberrima* additional chromosomal segments. Phenotypic effects of CSSL lines are represented in relative terms as circles on the right of the figure. The area of each circle is proportional to its relative effect. Horizontal bars at the bottom of the figure indicate QTL positions deduced from analyses performed using CSSL finder software. The colors of the circles and bars represent the effect as follows: red, increasing effect compared to the recurrent parent *Os_Caiapó*; green; decreasing effect.

**Table 2:**
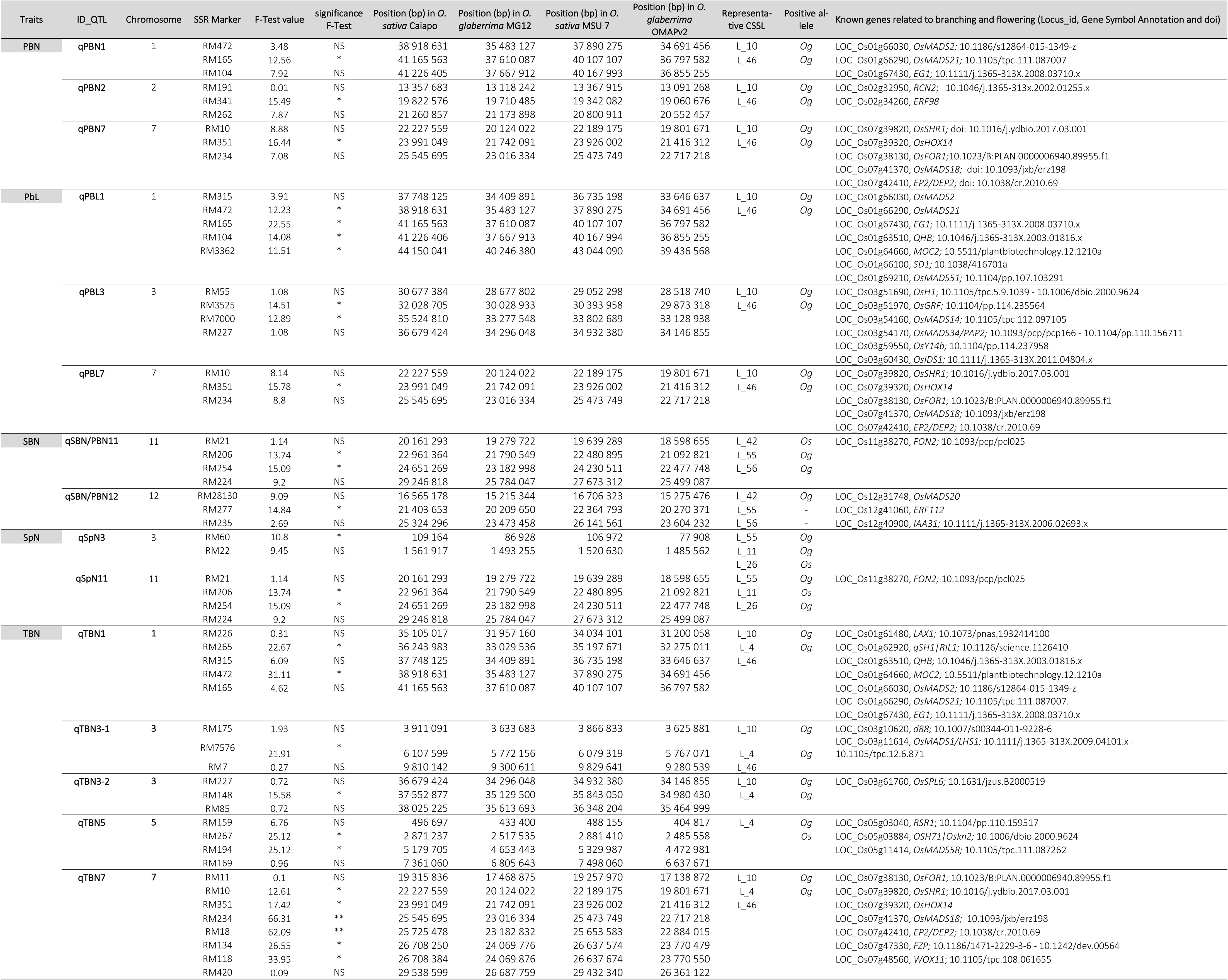
QTLs for morphological panicle traits detected in the CSSL population

*RL trait*: 13 CSSLs showed significant changes in RL value in comparison to the *Os_Caiapó* recurrent parent (Additional file1: Table S5). However, no QTLs were detected for this trait (Additional file2: Figure S1).

*SBN/PBN trait*: a total of 16 CSSLs showed a significant difference in secondary branch number per primary branch (SBN/PBN ratio) compared to the *Og_MG12* parent (Additional file 1: Table S5). Two QTLs associated with a decreased SBN/PBN ratio were detected (qSBN/PBN11 and qSBN/PBN12, maximum F-test scores of 15.09 and of 14.94 respectively) (Table 2; Additional file2: Figure S1). The average SBN/PBN ratio values were 3.0 for *Os_Caiapó* and of 1.7 for *Og_MG12*, corresponding to a decrease of 42% in *Og_MG12* relative to *Os_Caiapó*. The CSSLs L_42, L_55 and L_56 showed a decrease of 44.7%, 42.4% and 36.3% respectively relative to the *Os_Caiapó* parent, suggesting a high effect of the *Og_MG12* introgression(s) in these lines (Additional file1: TableS5). The phenotyping of these lines was repeated for a second year to confirm panicle trait variation (Additional file3: FigureS2). The L_55 and L_56 lines contain an *Og_MG12* segment in chromosome 11, but with missing marker information at the position of qSBN/PBN12. In contrast, the CSSL L_42 displays only an *Og_MG12* segment in chromosome 12 (Additional file3: Figure S2).

*SpN trait*: a total of 12 CSSLs showed a significant difference compared to the *Os_Caiapó* recurrent parent (Additional file 1: Table S5). Two QTLs (qSpN3 and qSpN11) were associated with a decreased SpN with maximum F-test scores of 10.8 and 15.1 respectively (Table 2; Additional file2: Figure S1). CSSLs L_55, L_11 and L_26 showed decreases of 37.9%, 32.3% and 30.8% respectively compared to the *Os_Caiapó* parent (Additional file1: Table S5). Lines L_55 and L_26 contain *Og* segments in both chromosomes 3 and 11 in the positions of the detected QTLs (Additional file3: Figure S2). Line L_11 contains only one *Og_MG12* introgression in chromosome 11, related to qSpN11 (Additional file 3: Figure S2).

*PBN trait*: a total of 13 CSSLs had significantly different PBN values compared to the *Os_Caiapó* recurrent parent (Additional file1: Table S5). Three QTLs corresponding to a decreased PBN (qPBN1, qPBN2 and qPBN7) with maximum F-test scores of 12.56, 15.49 and 16.44 were detected on chromosomes 1, 2 and 7 respectively (Table 2; Additional file2: Figure S1). Average PBN values were 11.9 for *Os_Caiapó* and 8.2 for *Og_MG12* (Additional file1: Table S5). The PBN values of two CSSLs (L_10 and L_46) differed significantly from the *Os_Caiapó* recurrent parent with a reduction of 24.1% and 20.9% respectively (Additional file 1: Table S5). These 2 CSSLs harbor similar *Og_MG12* segments in chromosomes 1, 2 and 7 (Additional file3: Figure S2).

*PBL trait*: Fourteen CSSLs exhibited significant differences with the *Os_Caiapó* recurrent parent for the PBL trait. Three QTLs corresponding to an increased PBL (qPBL1, qPBL3 and qPBL7) with maximum F-test scores of 22.55, 14.51 and 15.78 were detected on chromosomes 1, 3 and 7 respectively (Table 2; Additional file2: Figure S1). Average PBL values were 11.1 for *Os_Caiapó* and 12.3 for *Og_MG12* (Additional file1: Table S4). Two CSSLs (L_10 and L_46) showed increased PBL values of 19% and 12.8% respectively in comparison to the *Os_Caiapó* recurrent parent and harbored similar *Og_MG12* segments in chromosomes 1, 3 and 7 (Additional file1: Table S5; Additional file3: Figure S2).

*TBN trait*: Surprisingly, several CSSLs showed a significantly increased TBN value in our field conditions (Additional file1: Table S5) and five QTLs were detected: qTBN1, qTBN3-1, qTBN3-2, qTBN5 and qTBN7 with maximum F-test scores of 31.11, 21.91, 15.98, 25.12 and 66.09 respectively (Table 2; Additional file2: Figure S1). The L_10 and L_4 lines have similar *Og_MG12* segments in chromosomes 1, 3 and 7. Within chromosome 5, only the L_4 line contains a segment from the *Og_MG12* parent (Additional file3: Figure S2). As the L_46 line contains otherwise similar *Og_MG12* introgressions in chromosome 1, 3 and 7, this CSSL was included in the second phenotyping campaign, in which it revealed the presence of tertiary branches on its panicles in contrast to L_4 (Additional file3: Figure S2). Overall, these results support an association of *Og_MG12* introgressions with the presence of tertiary branches.

For the PBN, PBL and TBN panicle traits, the same BC_3_DH CSSLs (L_10 and L_46) led to the detection of several QTLs. The 2 lines in question display a similar panicle phenotype with a higher number of tertiary branches associated with a decreased PBN and increased PBL and SBN/PBN ratio values compared to the *Os_Caiapó* recurrent parent (Fig. 4A, B). The L_10 and L_46 BC_3_DH lines contain a complex association of *Og_MG12* introgressions in their genomes (Fig. 4C). For clarification, the regions corresponding to colocalized QTLs, associated with different traits, were renamed without specifying the associated traits: q_1 (for qPBL1, qPBN1 and qTBN1); q_2 (for qPBN2), q_3-1 (for qTBN3-1); q_3-2 (for qPBL3 and qTBN3-2); q_5 (for qTBN5); and q_7 (for qTBN7, qPBN7 and qPBL7).

**Fig. 4.**
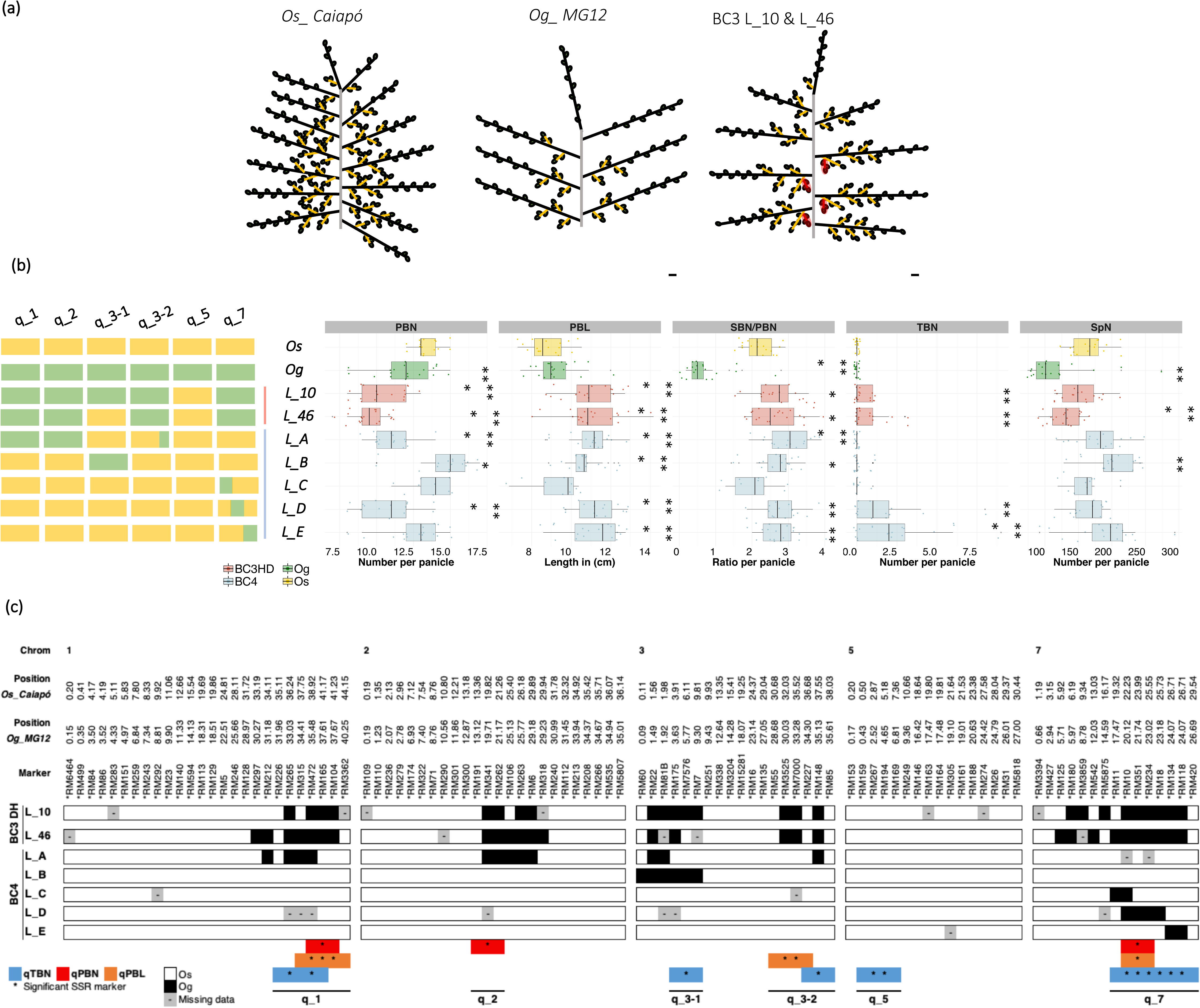
Phenotypic and genetic description of BC3DH and BC4 lines to dissect the effects of QTLs on panicle traits. **(a)** Schematic representation of the panicle structure observed in *Os_Caiapó*, *Og_MG12* and BC3DH L_10 and L_46, indicating primary branch (PB) in black, secondary branch (SB) in yellow, tertiary branch (TB) in red and spikelet (Sp) in green. Scale bar =1cm. **(b)** Boxplots of the phenotypic variation observed in Os_*Caiapó* (yellow), *Og_MG12* (green), BC3HD (red) and BC4 (blue) lines. Each point represents the phenotypic value for one panicle. Statistical significance (*t*-test *p*-values) between *Os_Caiapó* and each line for the panicle morphological traits is indicated as follows: ** *p*-values < 0.01; ***< 0.001. The left hand panel represents the allelic status (*Os_Caiapó* in yellow or *Og_MG12* in green) for each QTL region in each line. **(c)** Visualization of *Og_MG12* segments present in chromosomes 1, 3, 5 and 7 in BC3DH and BC4 lines to dissect the effects of QTLs on panicle traits. The positions of QTLs associated with PBN, PBL and TBN traits are indicated immediately above them (bars shaded in orange, red and blue respectively). Abbreviations: PBN, primary branch number; PBL, primary branch length; TBN, tertiary branch number; SBN/PBN, ratio between secondary branch and primary branch numbers; SpN, spikelet number.

### Substitutions of Os_Caiapó genomic regions by corresponding Og_MG12 segments can result in an added value panicle phenotype

As the altered phenotype observed in lines L_10 and L_46 was associated with multiple substitution segments (Fig. 4C), it was critical to determine whether this phenotype, and notably the formation of tertiary branches, involved one or several QTLs. Thus, different BC_4_F_3:5_ lines were obtained from the BC_3_DH lines by backcrossing and self-pollination (Fig. 4C), after which homozygosity was checked using the SSR markers carried by the introgressed *Og_MG12* segments present in the BC_3_DHs. Panicle traits were phenotyped for five BC_4_F_3_:_5_ lines containing the different individual *O. glaberrima* introgressed segments affecting TBN, together with *Os_Caiapó* and *Og_MG12* (Fig. 4B; Additional file1: Table S1). A second phenotypic campaign was carried out for validation of the BC_4_ lines L_B, L_D and L_E (Additional file4: Figure S3).

Divergent panicle traits compared to the *Os_Caiapó* parent were observed in all BC_4_ lines except for L_C (Fig. 5B). The latter contains an introgression at the beginning of q_7 (RM11 to RM10), suggesting that this region does not influence panicle architecture. All BC_4_ lines except L_C showed longer primary branches compared to the *Os_Caiapó* parent, suggesting that introgressions in chromosomes 1, 3 and 7 could independently influence primary branch length (Fig. 5A). The L_A line, containing *Og_MG12* regions corresponding to q_1, q_2 and q_3-2, also showed a decreased PBN value associated with a higher SBN/PBN ratio. However, no tertiary branches were observed. This suggests that the association of the q_1, q_2 and q_3-2 regions could induce a reduction in PBN associated with an increased SBN.

**Fig. 5.**
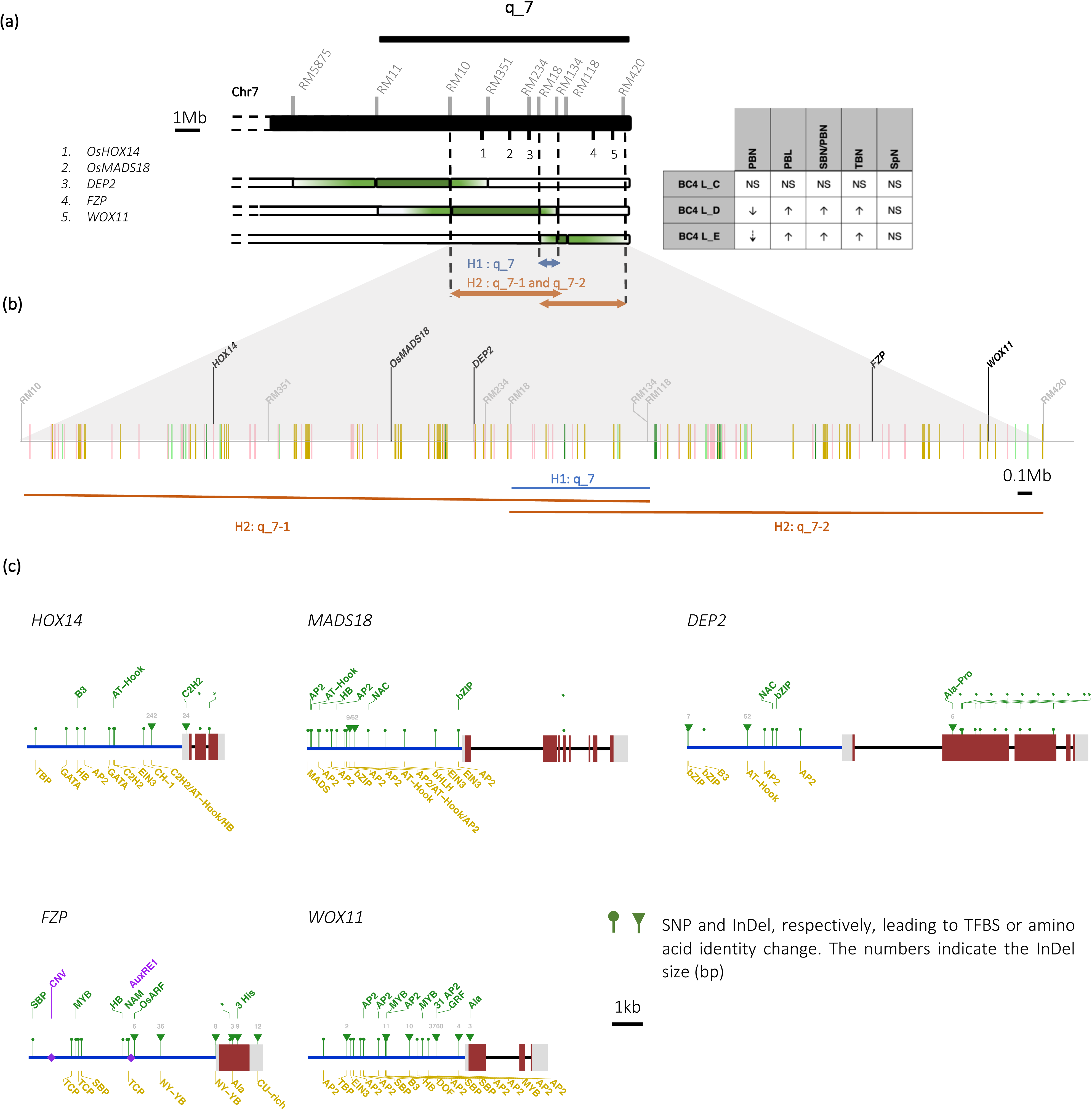
Description of genetic variations observed in the q_7 region between the *Os_Caiapó* and *Og_MG12* genomes **(a)** Influence of the q_7 region on panicle variation in the BC4 lines. Light green and green shading indicate the “extended” and strict introgression positions respectively for each BC4 line. **(b)** Representation of variation observed in the annotated genes between the *Os_Caiapó* and *Og_MG12* genomes in the region between the RM10 and RM420 SSR markers. Yellow bars show loci only present in *Os_Caiapó* genome, green bars indicate loci only present in *Og_MG12*. Light green bars represent genes that are duplicated in *Og_MG12* and pink bars represent genes that are annotated in *Og_MG12* but present in a different genomic region in *Os_Caiapó*. **(c)** Schematic representation of sequence divergence in candidate genes between the 2 genomes. Blue lines correspond to promoter regions, red and grey boxes represent exons and introns respectively. SNP and InDel variations leading to TFBS or amino acid changes are represented by dots and triangles respectively. The number above a triangle indicates InDel size. Asterisks indicate amino acid identity modification in the protein sequence. TFBSs present only in *Os_Caiapó* or *Og_MG12* are shown colored in yellow and green respectively. Abbreviations: TFBS, Transcription Factor Binding Site; Ala, alanine; Pro, proline; his, histidine; EIN3, Ethylene-Insensitive-Like3; C2H2, C2H2-Type Zinc finger; AP2, AP2-ERF; bHLH, basic helix-loop-helix; GRF, Growth-regulating factor; TCP, Teosinte-Branched1; SBN, Squamosa Binding Proetin; NY-YB, nuclear factor Y; HB, Homeodomain; MADS, MADS-BOX; bZIP, basic leucine zipper; TBP, TATA-box-binding.

Line L_B containing only one *Og_MG12* introgression (from RM60 to RM7) in place of q_3-1 showed increased values for four panicle traits (PBN, PBL, SBN and SpN) but not for TBN. However, these results were not observed during the second phenotyping of this line, except for an increase in PBL. Thus, it can be deduced that the q_3-1 region positively influences primary branch length in a manner that is independent of the other detected QTLs.

Finally, the BC_4_ lines L_D and L_E were the only ones observed to develop tertiary branches (Fig. 5A). In both lines, this phenotype is associated with an increased SBN/PBN and a longer PBL. The higher SBN/PBN ratio is associated with a decreased PBN in L_D, meaning that in this line more secondary branch meristems are established during panicle development. Similar results for SBN/PBN and TBN were observed in the second round of phenotyping, with a comparable decrease in PBN in the L_E line (Additional file4: FigureS3).

The two aforementioned lines contain contiguous introgression fragments in chromosome 7, from RM10 to RM18 for L_D and from RM134 to RM118 for L_E. Since the exact recombination positions of the introgressed segments for each BC_4_ is not known, the regions between SSRs with different genotypes are included in the QTL intervals. In Fig. 5A, it can be observed that lines L_D and L_E, which share the same phenotypes except for PBN, may harbor introgressions that overlap over only a small region between RM18 and RM134. Based on these observations, two different hypotheses can be proposed with regard to QTL position(s) in the q_7 region (Fig. 5A). The first one postulates the existence of a common QTL located between RM18 and RM134 that controls PBL, SBN/PBN and TBN. The second hypothesis is that two different QTLs, q_7-1 between RM10 and RM134, and q_7-2 between RM18 and RM420, exert a similar effect on PBL, SBN/PBN and TBN but act differently on PBN.

Taken together, the above results suggest that panicle architecture is a complex trait controlled by various different genomic regions which positively or negatively influence branching. An association of q_1, q_2 and q_3-2 produces opposing effects on PBN and SBN while only the q_3-1 region, comprised between RM175 and RM7, may impact positively upon PB length. Finally, the region in q_7, which negatively influences the number of primary branches and may be associated with the formation of tertiary branches and an increase in SBN per primary branch, may be defined as being either between the RM18 and RM134 marker positions or associated with two QTLs positioned between RM10 and RM134 for q_7-1 and from RM18 to RM420 for q_7-2.

### Co-location of QTLs with known genes and QTLs detected with other populations

Many studies have reported QTLs associated with panicle architecture and/or yield traits or both, using bi-parental (QTL mapping) or diversity panels GWAS (Crowell et al., 2014; Ya-fang et al., 2015; Bai et al., 2016b; Crowell et al., 2016; Rebolledo et al., 2016; Ta et al., 2018; Zhang et al., 2019; Cubry et al., 2020; Bai et al., 2021; Zhong et al., 2021). We found 70 common sites between the QTLs detected in this study and those reported earlier (Additional file 1: TableS6). For instance, co-located sites relating to similar traits (SBN and SpN) were found on chromosomes 11 and 12. Nevertheless, despite the significant number of common sites identified in the different studies, the traits that they have been reported to affect are not always strictly comparable or even similar, even if they relate in some way to panicle architecture or yield.

To evaluate the synteny of the QTL regions between the genomes of the two CSSL parents, sequences from the assemblies of the *Os_Caiapó* and *Og_MG12* genomes obtained by ONT sequencing (https://doi.org/10.23708/QMM2WH; https://doi.org/10.23708/1JST4X; Additional file1: TableS7) were used to carry out local alignment for each QTL region. The assembled genomes are of high continuity and completeness, with BUSCO score of 98.4% allowing precise detection of structural variations. For the majority of the QTL regions, no major structural variation was observed between the two genomes (Additional file5: Figure S4). However, some notable differences were detected at the qPBN2, qTBN5, qTBN7, qSpN3 and qSBN12/PBN12 sites. Within the QTLs qPBN2 and qTBN7, inverted segments of 1,296,899 and 230,762 bp were observed between *Os_Caiapó* and *Og_MG12* respectively (Additional file1: Table S8). The genomic regions for QTLs qTBN5, qSpN3 and qSBN/PBN12 displayed InDels between *Os_Caiapó* and *Og_MG12* (Additional file1: Table S8). With the exception of qSBN11, we observed a high sequence similarity in the QTL regions between the two genomes, suggesting similar content in terms of coding sequences. Synteny analysis was extended by the alignment of the *Os_Caiapó* and *Og_MG12* QTL regions with those of the *O. sativa* cv. Nipponbare reference genome (Additional file5: Figure S4). Based on the high conservation of the corresponding regions between *Os_Caiapó* and *Os_Nipponbare*, we then used as a reference, for subsequent analyses, the *O. sativa* cv. Nipponbare (IRGSPv1.0) functional gene annotation databases (i.e., MSU7 and RAP_db) for candidate gene prioritization in relation to panicle architecture and flowering.

For all QTLs identified in the present study, we found several known genes with relevant functions that related to flowering and/or panicle development (Table 2; Additional file6: Figure S5). Genes such as *LAX PANICLE1* (*LAX1*), *OsMADS1/LEAFY HULL STERILE1* (*LHS1*), *OsMADS14*, *OsMADS34/PANICLE PHYTOMER2* (*PAP2*), *OsINDETERMINATE SPIKELET 1 (OsIDS1*), *OsMADS18*, *DENSE AND ERECT PANICLE2* (*DEP2*) and *FRIZZY PANICLE* (*FZP*) are known to be involved in the control of the panicle development and were suggested as candidates that might contribute to the panicle branching diversity observed between the two parents *Os_Caiapó* and *Og_MG12* (Chongloi et al., 2019; Wang et al., 2021; Yin et al., 2021).

### q_7 genetic variations between Os_Caiapó and Og_MG12

We paid particular attention to the q_7 region, as it was found to be associated with several panicle morphological traits and to span a genomic region that included several panicle-associated genes. To understand if specific genetic modifications present in the q_7 QTL region could be associated with panicle architecture variation, we further compared the constituent genes in the region between the RM10 and RM420 markers in the *Os_Caiapó* and *Og_MG12* genomes on chromosome 7 (22.23 – 29.59 Mbp in *Os_Caiapó* vs 20.12 – 26.69 Mbp in *Og_MG12*). For this purpose, gene annotation comparison and BLAST analyses were performed to explore in detail gene synteny and presence/absence of genes between the *Os_Caiapó* and *Og_MG12* genomes within this region (Fig. 5B; Additional file1: Table S9). Between the RM10 and RM420 marker positions, we observed a variation in the number of genes between the two genomes (Additional file1: Table S9). This variation is due to several factors: the differential presence of corresponding orthologs between the two genomes; differential gene duplication within the region; and different locations of a given gene within the two genomes (i.e., genomic rearrangement) (Fig. 5B; Additional file1: Table S9). Among the genes that differ between the two genomes in this region, many are related to transposable elements (TEs) or are hypothetical protein-encoding genes. None of them has a known function relating to panicle development or flowering control, nor to any other developmental processes.

Special attention was paid to common genes present in the q_7 region that were known to be related to inflorescence development and/or meristem activity and maintenance. Based on a bibliographic survey, no candidate genes of special interest were identified within the region between the RM18 and RM134 markers. In contrast, several interesting candidate genes were identified in the q_7-1 and q_7-2 regions: the *OsHOX14*, *OsMADS18* and *DEP2* genes in q_7-1 and the *FZP* and *WOX11* genes in q_7-2 (Fig. 5C). SNP and InDel sites associated with the aforementioned genes that were polymorphic between the *Os_Caiapó* and *Og_MG12* genomes were analyzed in order to detect amino acid modifications, open reading frame alterations or transcription factor binding site (TFBS) variations within promoter regions.

*OsHOX14* (LOC_Os07g39320/Os07g0581700) encodes a protein of the Homeodomain-leucine zipper (HD-Zip) TF family and is involved in the regulation of panicle development (Beretta et al., 2023; Shao et al., 2018). Two of the identified SNPs cause amino acid variations in the coding sequence, including one in the homeobox domain (Fig. 5C; Additional file1: Table S10). Comparison of the *OsHOX14* and *OgHOX14* promoter regions revealed InDel and SNP variations leading to the loss of 9 TFBSs and a gain of three new TFBSs in the CSSLs bearing *Og_MG12* q_7-1 introgression (Fig. 5C; Additional file1: Table S11).

*OsMADS18* (LOC_Os07g41370/Os07g0605200) encodes a protein of the AP1/FUL-like MADS-box TF subfamily (Masiero et al., 2002). In addition to its role in flowering time promotion (Fornara et al., 2004), *OsMADS18* is known to specify inflorescence meristem identity (Kobayashi et al., 2012). SNPs were detected in the coding sequence; one of the SNPs leads to a non-synonymous change in the OgMADS18 protein outside the known binding or functional domains (Fig. 5C; Additional file 1: Table S10). Scanning of the promoter regions of *OsMADS18* and *OgMADS18* in the *Os_Caiapó* and *Og_MG12* genomes respectively revealed several SNPs and InDels that lead to variations between the two genomes (presence/absence) of TFBSs recognized by AP2, NAC or HB TFs (Fig. 5C, Additional file1: Table S11).

*DEP2* (LOC_Os07g42410/Os07g061600) encodes a plant-specific protein of unknown function associated with panicle size (Li et al., 2010). Various SNPs and one InDel are observed between the coding sequences of *OsDEP2* and *OgDEP2* which result respectively in changed amino acid identities between the two protein orthologs (Fig. 5C; Additional file 1: Table S10) and an insertion of two amino acids in the OgDEP2 protein. Sequence variations were also observed between the promoters of *OsDEP2* and *OgDEP2*; these variations lead to the loss of bZIP, B3, AT-Hook and AP2 TFBSs in the CSSLs carrying q_7. On the other hand, in comparison to *O. sativa*, the CSSLs carrying q_7 include NAC and bZIP TFBSs specific to the promoter of *OgDEP2*.

Moreover, among the genes contained in the q_7-2 region, *FZP* (LOC_Os07g47330/Os07g0669500) encodes an AP2/ERF TF known to play a role in the regulation of floral meristem determinacy in rice (Komatsu et al., 2003; Zhu et al., 2003). Several SNP and InDel variations observed in the coding region of *FZP* lead to coding sequence amino acid changes between the two genomes (Fig. 5C; Additional file 1: Table S10). An InDel of 9 bp in *OgFZP* results in the insertion of three histidine amino acids outside the single AP2 domain in the OgFZP protein. Several recent studies have shown that variations in the promoter region and 3’ UTR (Untranslated Transcribed Region) of the *OsFZP* gene affect panicle architecture by through changes in the number of branches produced (Bai et al., 2016, 2017; Huang et al., 2018; Wang et al., 2020; Chen et al., 2022). Here, we observed several SNPs and InDels between the promoter of the two orthologs, which result in a differential presence of TFBSs (Fig. 5C, Additional file1: Table S11). No variation was observed in the copy number variant (CNV) motif of 18 bp described by Bai et al. (2017). Huang et al. (2018) revealed a deletion of 4 bp in the *OsFZP* 5’ regulatory region in comparison to the sequence of *O. rufipogon*. This deletion is observed in *Os_Caiapó* and not in the genome of *Og_MG12* (Fig. 5C). Recently, it has been shown that CU-rich elements (CUREs) present in the 3’ UTR of the *OsFZP* mRNA are crucial for efficient OsFZP translational repression (Chen et al., 2022). Three CUREs are detected in the 3’ UTR of *OsFZP* in the *Os_Caiapó* genome. In contrast, a deletion of the third CURE sequence is observed in the *Og_MG12* genome (Fig. 5C).

Although the WUSCHEL-related homeobox *WOX11* gene (LOC_Os07g48560/Os07g0684900) has been mainly described as playing an important role in root development (Zhao et al., 2009; Zhang et al., 2018), this gene is also required to regulate rice shoot development and panicle architecture (Cheng et al., 2018). An InDel in the coding region of *WOX11* leads to an Asn amino acid insertion in the OgWOX11 protein in *Og_MG12* without affecting the homeobox domain of the protein (Fig. 5C, Additional file1: Table S10). In the promoter region, variations between *Os_Caiapó* and *Og_MG12* lead to the presence/absence of several TFBSs in the promoter of *OgWOX11* (Fig. 5C; Additional file 1: Table S11). We also observed a large insertion of 3,760 bp, containing about 31 AP2 TFBs, in *Og_MG12* compared to the *Os_Caiapó* genome.

Overall, our analysis revealed variations in synteny that demonstrate an absence of certain *Os_Caiapó* genes and the addition of other loci as a consequence of *Og_MG12* genomic introgressions within the CSSLs: some of these changes can be hypothesized to play a role in determining panicle trait diversity. Moreover, our analysis of candidate genes revealed TFBS variations and protein coding sequence polymorphisms that may lead to variations in transcript levels and/or protein activity in CSSLs harboring the q_7 introgression.

## Discussion

### Added value of the O. sativa x O. glaberrima CSSL population

Inter-specific *O. sativa* x *O. glaberrima* CSSL genetic resources are of great interest, as *O. glaberrima* can provide a gene pool with potential for rice improvement in terms of resistance to biotic and abiotic stresses and ecological adaptability (Gutiérrez et al. 2010). CSSL libraries are also good pre-breeding materials for the simultaneous identification, transfer and pyramiding of key genes in crop improvement programs. They are also useful for the study of traits lost or retained during the process of evolution, domestication and selection (Ali et al., 2010; Balakrishnan et al., 2019). Genetic strategies based on CSSL populations have the advantage of using a relatively small number of lines for experiments, leading to the possibility of replicating evaluations, thereby enhancing statistical strength for complex and time-consuming phenotypic assessments.

In this study, backcross introgression lines (BC_3_DH) of *O. sativa_Caiapó* x *O. glaberrima_MG12* were phenotyped for panicle morphological traits, which are key components of yield potential. Our analyses led to the detection of 15 QTLs, localized on chromosomes 1, 2, 3, 5 and 7 and comprising several genes known to be involved in panicle architecture determination (Table 1). Colocalization was observed with previous QTLs and GWAS sites associated with panicle branching traits, supporting the implication of the QTLs detected in this study in the regulation of panicle morphology. Moreover, since the genomic sites described here were detected using various different sets of populations, we propose that the genomic regions in question play a major role in determination of panicle architecture diversity.

Some of the BC_3_DH lines used in this analysis contain several *Og_MG12* introgression regions, thus potentially allowing the detection of several QTLs with epistatic interactions. For most of the traits studied, we detected several QTLs within the same line, as observed in the lines L_10 and L_46 that allowed the localization of QTLs governing TBN-, PBN- and PBL-related traits. This is not surprising in the case of a complex trait such as panicle architecture, which is controlled by a large number of genes and QTLs with small effects that could be influenced by environmental and epistatic interactions (Xing et al., 2002). An additional advantage of the CSSL-based approach is that it provides novel interspecific combinations of parental alleles which may significantly affect panicle branching traits, thereby providing an insight into how inflorescence architecture is governed in the two species and how their respective genetic factors interact. The fact that we observed a colocalization of QTLs for different panicle traits suggests a likely genetic interdependence between these traits; this hypothesis is supported by correlations observed between phenotypic values for morphological characters such as SpN, SBN, PBN, PBL and RL that share common QTLs.

### Impact of O. glaberrima introgressions on panicle traits in the O. sativa background

Compared to the *Os_Caiapó* recurrent parent, the *Og_MG12* donor parent displays a global reduction in panicle constituent numbers (i.e., numbers of primary branches, secondary branches and spikelets) and produces longer primary branches. Tertiary branches were not observed in the *Og_MG12* donor parent. On the basis of these observations, the QTLs detected in the CSSLs were expected to be associated with a global decrease in panicle constituent numbers (PBN, SBN, SpN) and an increase of PB length in comparison to the *Os_Caiapó* parent. Indeed, this is mainly what was observed. In most lines studied, introgressions of *Og_MG12* alleles of panicle-regulating genes produced a negative effect on the regulatory network controlling panicle morphological traits, indicating that some of the genomic variations observed between *Og_MG12* and *Os_Caiapó* in these regions are functionally important.

Surprisingly, we also detected QTLs associated with an increase in panicle trait values in comparison to the *Os_Caiapó* recurrent parent. Two of the BC_3_DH CSSLs (L_10 and L_46) carried four substitution segments from the *Og_MG12* parent donor and displayed panicle branching modifications that included not only a reduction of PBN and an increase of PBL as expected, but also an increase in secondary branch number per primary branch and a formation and development of tertiary branches (TB). As confirmed with the corresponding BC_4_ lines, both of the CSSL lines displayed a phenotype with an added value for one or several panicle traits. For example, the *Og_MG12* introgression localized on chromosome 3 in place of q_3-1 induces a globally higher complexity of panicle branching with an increase in PBN, SBN/PBN and SpN, associated with an increased PBL. This region includes the gene *OsMADS1/LHS1* belonging to the *SEPALLATA MADS-Box* TF family which is known to play an important role in determining floral meristem identity and in floral organ development (Jeon et al., 2000; Prasad et al., 2005; Cui et al., 2010). In *O. sativa*, *OsMADS1/LHS1* directly regulates several other transcription factor genes (MADS box and Homeodomain family members) and hormone signaling pathways implicated in floral meristem specification, maintenance and determinacy (Khanday et al., 2013).

The unexpected results obtained for the aforementioned lines indicate that introgression of certain *Og_MG12* regions into the *Os_Caiapó* genetic background can lead to an enhanced panicle phenotype and suggest a perturbation of the pathways regulating inflorescence development in *Os_Caiapó*. This perturbation could be explained by several mechanisms. Firstly, it is possible to postulate the presence of a negative regulator of branching in *Os_Caiapó* that is absent in the *Og_MG12* introgression, leading to an upregulation of branching in the CSSL. A second hypothesis could be proposed whereby one or more genes specific to *O*. *glaberrima* are present in the *Og_MG12* introgression compared to the corresponding *Os_Caiapó* region. One or more of these genes could be positive regulators of panicle architecture, with this regulatory potential being either latent or only very weakly expressed in the *Og_MG12* background. A third possible explanation could be proposed involving *cis*-variations associated with orthologous genes conserved between the two genomes. In this case, the addition of new *Og_MG12* genes, or their variant allelic forms into the *Os_Caiapó* genomic background could be postulated to exert an additive effect on panicle branching in the CSSLs, by affecting regulatory interactions within the *Os_Caiapó* network governing panicle branching.

In a similar way, our observations on the effects of QTL q_7, which favored increased SB and TB numbers, might have more than one possible explanation. Comparative analysis of the BC_4_ lines led us to suggest two hypotheses (above) regarding the position of QTL(s) on q_7 region (Fig. 5A). However, based on quantitative and qualitative differences between the BC_4_ line phenotypes, we favor the hypothesis whereby the presence of two distinct regions could influence panicle trait variation differentially (i.e., the q_7-1 and q_7-2 regions). In this scenario, both would be associated with increased PBL, SBN and TBN traits values whereas q_7-1 would have an additional negative effect on PBN. Moreover, the overlapping region of the introgressed segments in lines L_D and L_E, which spans a region of ∼984 Kbp, is recombined in both lines. This implies that Lines L_D and L_E can bear the *Os* allele at any locus in this region so the the probability that both lines bear the *Og* allele at a same putative QTL position is low. In addition, this region only contains a few genes whose predicted function is unknown.

In parallel, we observed a good synteny between the two species within the q_7-1 and q_7-2 regions, ruling out the possibility that large-scale genomic rearrangements might explain the phenotypic variations observed. Indeed, the strong global synteny between these regions suggests rather that CSSL phenotypes might be accounted for by modifications within the genomic regions of key regulatory genes that give rise to variations in their expression or in their protein functions and/or interactions through gene regulatory networks (GRNs). Mutations affecting transcription factor proteins and/or DNA sequences or both, such as enhancers and promoters, fall into this category (Hill et al., 2021). Moreover, traits associated with dynamic processes are more readily modified through their *cis*- and/or *trans*-regulation rather than through coding sequence mutations (Wray, 2007). The q_7-1 and q_7-2 regions comprise several genes known for their role in panicle development and numerous SNPs and InDels have been detected in the promoter regions and coding sequences of these candidate genes.

Within the q_7-1 region, three genes have been reported to have a direct function in panicle architecture establishment. The homeodomain-leucine zipper transcription factor gene *OsHOX14* has been shown to be involved in the regulation of panicle development and its loss of function leads to a reduction in PBN compared to wild type (Shao et al., 2018; Beretta et al., 2023). The second candidate gene *OsMADS18*, in addition to its role in flowering time promotion (Fornara et al., 2004), is known to specify inflorescence meristem identity through interaction with PANICLE PHYTOMER2 (PAP2/OsMADS34) and the two other members of the AP1/FUL subfamily, OsMADS14 and OsMADS15 (Kobayashi et al., 2012). *DEP2* encodes a plant-specific protein without a known functional domain, mutations of which affect elongation of the rachis and both primary and secondary branches, caused by a defect in cell proliferation (Li et al., 2010; Mao et al., 2020; Wang et al., 2021b). For each of these three genes, the *Og_MG12* allelic version shows variations in comparison to *Os_Caiapó* in both protein coding sequences and in putative TFBSs within the promoter region. The presence of these binding sites may reveal a divergence in the regulation of the expression of these genes between the two parents which could lead to the morphological variations observed in the CSSLs (i.e., decrease in PBN).

Concerning the q_7-2 region associated with the formation of higher order panicle branches (i.e., secondary and tertiary branches), the candidate genes *WOX11* and *FZP* were detected in both genomes. The WUSCHEL-related homeobox gene *OsWOX11* is necessary and sufficient to promote crown root emergence and growth in rice by responding to or regulating auxin and cytokinin signaling (Zhao et al., 2009, 2015; Cheng et al., 2014, 2015; Zhang et al., 2018). OsWOX11 can recruit a histone aceyltransferase complex to activate downstream target genes to stimulate cell proliferation in crown root development (Zhou et al., 2017). Cheng et al. (2018) showed that OsWOX11 and JMJ705 cooperatively control shoot growth and commonly regulate the expression of a set of genes involved in meristem identity (Cheng et al., 2018) indicating that the role of this gene is not confined to roots. A comparison of the promoter regions of *WOX11* orthologs revealed numerous SNP and InDel differences between the two genomes and an insertion of 3,760 bp in *Og_MG12*, which includes numerous AP2 TFBS potentially recognized by the *MULTI-FLORET SPIKELET 1 (MFS1)* and *OsFZP* genes, known to play an important role in the regulation of spikelet meristem determinacy (Komatsu et al., 2001, 2003; Zhu et al., 2003; Ren et al., 2013). In an *O. sativa* genomic environment, the promoter of *OgWOX11* containing new AP2 TFBSs could potentially be programmed differently in its expression during the reproductive phase of the plant due to altered promoter activity, with possibly effects on cell proliferation in the axillary meristem leading to higher order branching in the CSSL lines.

The *OsFZP* gene plays an important role for the transition from branch to spikelet primordium during panicle development (Komatsu et al., 2001, 2003; Zhu et al., 2003). Reduced expression of *OsFZP* at the reproductive stage increases the extent of higher order branching of the panicle, resulting in increased grain number (Bai et al., 2016). Moreover, recent studies revealed *OsFZP* as the causal gene of QTLs for grain number per panicle (Bai et al., 2017; Fujishiro et al., 2018; Huang et al., 2018; Wang et al., 2020) and showed that the fine-tuning of *OsFZP* expression at the transcriptional and post-translational levels could determine panicle architecture (Bai et al., 2017; Huang et al., 2018; Chen et al., 2022). Some SNPs and InDel variations between the *Os_Caiapó* and *Og_MG12* genomes were detected within the promoter region of the two *FZP* orthologs. Notably, the *OgFZP* promoter does not possess a 4bp deletion observed in the *Os_Caiapó* genome adjacent to an auxin response element (AuxRE) 2.7 Kbp upstream *OsFZP*. This deletion is known to affect the binding activity of OsARF6 to the promoter of *OsFZP*, leading to decreased accumulation of *OsFZP* mRNAs, in comparison to its *O. rufipogon* Griff. wild-relative ortholog. This reduced amount is associated with an increase in secondary branch number in *O. sativa*, in comparison to *O. rufipogon* (Huang et al., 2018). Consequently, *OgFZP* is more similar to its *O. rufipogon* ortholog in this part of the regulatory region. We also observed a deletion in the CU-rich elements (CUREs) localized in the 3’UTR of *OgFZP* in comparison to its *Og_Caiapó* ortholog. This element is known to be crucial for efficient OsFZP translation repression, via the binding with OsPTB1 and OsPTB2 polypyrimidine trans-binding proteins acting as repressors (Chen et al., 2022). Accordingly, these two variations observed in regulatory regions of *OgFZP* could result in an increase in transcript and/or protein amounts, leading to a precocious transition from branch to spikelet primordium. Such a scenario would be corroborated by the less-branched panicle phenotype of *Og_MG12* compared to *Os_Caiapó* but not by the phenotype of the CSSLs containing *OgFZP*. The analysis of the combinatorial effect of variations in these regulatory elements would be useful to explain the observed phenotypes.

## Conclusion

This study was carried out with the broad aim of identifying discrete genetic elements that govern rice panicle architectural diversity using a population of interspecific introgression lines. We detected several QTLs associated with rice yield and panicle branching diversity observed between the two cultivated rice species *O. sativa* and *O. glaberrima*, with decreased values – as expected – in the CSSLs but also unexpectedly with added values for some QTLs compared to the *O. sativa* recurrent parent. This was the case for QTL q_7 on chromosome 7, which causes a global modification of panicle morphology due to two distinct genomic regions: q_7-1 and q_7-2. For both regions, the *Og_MG12* introgression causes an increase in higher orders of branches and q_7-1 is additionally associated with a reduction in PBN. A detailed comparative genomic analysis revealed variations in gene content, promoter region sequence and protein coding sequences for a number of different key genes that act in the control of panicle architecture. When an *Og_MG12* allelic variant is placed in an *Os_Caiapó* genomic background context, it might have an additive effect on panicle branching in the CSSLs, through its addition to the *Os_Caiapó* gene regulatory network (GRN) regulating panicle branching. Moreover, it cannot be ruled out that the higher complexity panicle architecture observed in corresponding CSSLs could result from the combination of these key genes in the *Os_Caiapó* GRN. However, it should be borne in mind that the biological functions and regulation of the genes present in the q_7-1 and q_7-2 regions have been described in *O. sativa*. The functional analysis of their homologs in *O. glaberrima* has yet to be investigated to confirm similar roles and/or to assess their regulation. Further analysis of the regulation of these genes will be beneficial in improving rice breeding for the exploitation of favorable alleles present in this CSSL population. Some genomic regions associated with multiple traits point to the possibility of enhancing yield potential. The genetic material created here will provide knowledge and resources to facilitate the retention of favorable alleles in breeding programs and to develop new improved rice cultivars using *O. glaberrima* as a genetic resource for enhancing morphological traits in *O. sativa*.

## Materials and Methods

### Plant Materials

For QTL detection, a set of 60 introgression lines, or Chromosome Segment Substitution Lines (CSSL) was used. The CSSLs were derived from a cross between *O. sativa* subgroup tropical japonica (cv. Caiapó) as the recurrent parent and *O. glaberrima* (cv. MG12; acc. IRGC103544) as the donor parent (Gutiérrez et al., 2010). The CSSLs used were BC_3_DH, that is, obtained after three rounds of backcrossing to the recurrent parent followed by double haploidization of BC_3_F_1_ male gametes (Additional file 1: Table S1). The selection of this set of 60 lines covering the entire donor genome from an initial panel of 312 BC_3_DH lines genotyped using 200 simple-sequence repeat (SSR) markers was carried out with the help of the program CSSL Finder v0.9 (http://mapdisto.free.fr; Gutiérrez et al., 2010). The effects of *O. glaberrima* introgressions on panicle trait variation were evaluated by choosing five BC_4_F_3:5_ lines, each containing the target region and as few introgressed *O. glaberrima* genomic segments in the rest of the genome as possible (Additional file 1: Table S1).

### Phenotypic evaluation

The 60 BC_3_DH lines, along with the parents (*Os*_*Caiapó* and *Og*_*MG12*), were grown together in November 2012-January 2013 (Year 1) in the International Center for Tropical Agriculture (CIAT, now Alliance Bioversity-CIAT) headquarters experimental fields (Palmira, Colombia) (76° 21’W, 3° 30’N and 967masl) under irrigated conditions. The experimental design was a randomized complete block design with two replications and 62 plots. Fives plants per line per plot were grown. Panicle traits were evaluated for subsequent QTL analysis and validation. For each line, the three main panicles from three randomly chosen plants (9 panicles total) per line per replicate were collected. Each panicle was spread out on a white background and held in place with metal pins for photography shooting. A total of 1,443 panicles from the 60 BC_3_DH lines and the parents were dissected and scored manually. In January-April 2014 (Year 2), a total of eight BC_3_DH lines were re-evaluated using P-TRAP software (Al-Tam et al., 2013), which performs automatic scoring of panicle traits. Five BC_4_F_3_/F_5_ lines with parents, plus L_10 and L_46 BC_3_DH, were grown in the greenhouse in Montpellier, France (March 2021) at 28°C / 80% relative humidity under short days conditions (11h light-13h dark), and were scored for panicle traits using P-TRAP. Panicle phenotypes were re-evaluated in the same conditions for three BC_4_ lines (June 2021).

For each analysis (2013, 2014 and 2021), 6 morphological panicle traits were scored: rachis length (RL); number of primary branches (PBN); number of secondary branches (SBN); number of tertiary branches (TBN); spikelet number (SpN); and average primary branch length (PBL) per panicle (Additional file 1: Table S2).

### Statistical Analysis

Basic statistical analyses were performed for each trait using Analysis of variance (ANOVA) functions in the R software package (version 4.0.4, R Core Team 2022), taking into consideration all factors (replicates, blocks within replicates and lines) as fixed effects. Trait heritability was computed using the line effect based on the variance among phenotypic measurements between the two replicates of the population phenotyping assay. The corrplot, ade4 and devtool R packages were used to analyze phenotypic correlation between traits.

The CSSL Finder program was used to display graphical genotypes of the 200 SSRs in the 60 BC_3_DH CSSLs together with phenotypic values for each trait. QTLs were detected using the graphical genotyping tool proposed by CSSL Finder, that allows combination of logical genotype-phenotype association with single-marker ANOVA1 F-test. The F-test was considered significant when its value was higher than 10.0.

Comparison between the CSSLs and the recurrent parent *Os*_*Caiapó* was performed by a Dunnett’s multiple comparison test (! < 0.05). The relative effect *RE* of an *O. glaberrima* introgression in a given CSSL was calculated using the least square means (LSMEANS) output of the GLM procedure as follows:

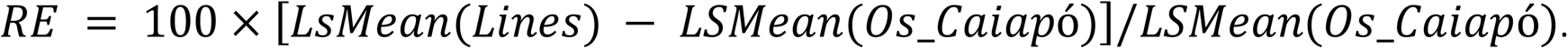

A QTL was taken into consideration when the trait value of a given line was significantly different from that of the parent *Os_Caiapó* (! < 0.001) and when at least two lines having overlapping segments shared similar phenotypes. The phenotypic variation of these lines was further evaluated in a second phenotyping experiment in Year 2.

### Genome sequencing, assembly, quality assessment and annotation

Samples of young leaves from *O. sativa Caiapó* and *O. glaberrima MG12* were used to extract genomic DNA and for preparation of SKL-LSK109 libraries as described in Serret et al., 2021.

Genome sequencing was performed on a Nanopore MinION Flow Cell R9.4.1 (Oxford technologies, Oxford Science Park, UK). The long-read sequences of *Os_Caiapó* and *Og_MG12* generated by the ONT sequencing were assembled using the following steps. After basecalling and Q>8 filtering with Guppy V6.1.2, with the SUP model, the preliminary genome was assembled with Flye V2.9 and default assembly parameters for Sup ONT (Kolmogorov et al. 2019). Medaka V1.6.1 was then used to create consensus sequences and to improve the accuracy of the assembly (polishing). The contigs were ordered with RagTag V2.1.0 using *O.sativa cv Nipponbare/IRGSP1.0* as a reference and with default parameters (Alonge et al., 2022). An evaluation of genome assembly completeness was carried out using BUSCO (Benchmark Universal Single Copy Orthologs) v5.2.2 with default parameters and the Poale database (Additional file1: Table S7) (Manni et al., 2021).

The Liftoff tool was used for gene prediction and annotation using the genome reference *O. sativa* cv Nipponbare (IRGSPv1.0, MSU7.0) and associated gff files for both *Os_Caiapó* and *Os_MG12* and by using the *O. glaberrima* genome reference OglaRS2 along with its associated gff files (https://ftp.gramene.org/oryza/release-6/gff3/oryza_glaberrima/) in the case of *Og_MG12* (Additional file1: Table S7) (Tranchant-Dubreuil et al., 2023).

### Genomic Alignment

The sequences of the QTL regions and the genes of interest were extracted from the three genomes *O. sativa* cv. *Caiapó*, *O. glaberrima* cv. *MG12* and *O. sativa* cv. *Nipponbare* MSU7 using Seqkit version 2.4.0 (Shen et al. 2016). Sequences of QTL regions were aligned and plotted using the minimap2 aligner implemented in the D-GENIES online tool (https://dgenies.toulouse.inra.fr/run) using the option Few repeats).

### QTL colocalization, identification and bio-analysis of candidate genes

For QTL and gene colocalization, the qTARO database (http://qtaro.abr.affrc.go.jp), the funRiceGenes database (https://funricegenes.github.io/) (Huang et al. 2022) and various datasets from recently published works were used to identify QTLs and genes that overlapped with the QTL regions detected in this study. To identify genes potentially associated with the QTL regions, we used the *O. sativa* cv *Nipponbare* MSU7.0 (https://riceplantbiology.msu.edu) and the RAP_db (https://rapdb.dna.affrc.go.jp) annotation databases. From the annotated gene list, the candidate genes were identified based on their predicted function (biological processes), their referencing in the funRiceGenes database and/or their expression pattern with respect to the trait of interest (Harrop et al., 2016; Harrop et al., 2019). Regions spanning the candidate genes and their associated 5 Kbp upstream region (considered as their promoter regions) were obtained for *O. sativa* (IRGSPv1.0, MSU7.0) and for *O. glaberrima* (OglaRS2), then nucleotide and protein alignments were performed using CLUSTALW in order to identify SNPs, InDels and amino acid changes between the two species (Thompson et al. 1994). The Plant Promoter Analysis Navigator web facility (http://plantpan.itps.ncku.edu.tw/) was used to detect transcription factor binding sites (TFBSs) and regulatory elements (CpG islands and tandem repeats), with a cut-off at 0.8, in the promoter region of candidate genes (Chow et al., 2018). Variations between the two genomes, in terms of promoter and gene structures along with SNP and InDel positions and information, were drawn using the R package KaryoploteR (Gel and Serra 2017).

## Abbreviations

RL: Rachis Length
PBN: Primary Branch Number
PBL: Primary Branch Length
SBN: Secondary Branch Number
SpN: Spikelet Number
SBN/PBN: ratio of SBN and PBN
*Os_Caiapó*: *Oryza sativa* cv. *Caiapó*
*Og_MG12*: *Oryza glaberrima cv. MG12*
CSSL: Chromosome Segment Substitution Lines
BC: backcross
SRR: Simple Sequence Repeat
TFBS: Transcription factor Binding Site.

## Acknowledgments

The authors acknowledge Ndomassi Tando and the ISO 9001 certified IRD itrop HPC (member of the South Green Platform) at IRD Montpellier for providing HPC resources that have contributed to the research results reported within this paper (URL: https://bioinfo.ird.fr/-http://www.southgreen.fr). We thank Lady Johanna Arbelaez Rivera (Alliance Bioversity-CIAT), Sophie Chéron (IRD) and Harold Chrestin (IRD) for plant care. We thank Marie-Christine Combes (IRD) and Carole Gauron (IRD) for their help with the panicle phenotyping work. We thank Jean-Francois Rami (Cirad) for his help with R scripts. We also thank Laurence Albar (IRD) for her helpful feedback on the manuscript.

## Funding

This research was funded by the Agropolis Foundation through the “Investissements d’avenir” programme (ANR-10-LABX-0001-01), the Fondazione Cariplo (EVOREPRICE 1201-004), and the CGIAR Research program on Rice (RICE CRP).

## Availability of data and materials

The CSSLs materials are available from the corresponding authors on reasonable request. Sequencing data generated for this project are available at https://doi.org/10.23708/QMM2WH and https://doi.org/10.23708/1JST4X for *Os_Caiapó* and *Og_MG12* respectively.

## Authors’ Contributions

HA, ML designed the project, with input from SJ; HA and ML performed QTL analysis; ML designed the population subset and the markers and supervised field phenotyping; AG performed the SSR analysis on the BC_3_DH and BC_4_ plants. MC and JS performed the ONT sequencing. MC and FS performed genome assembly and annotation transfer. FN performed the bioinformatic minimap2 alignments. JO developed a Python script for the colocalization analysis. HA, SJ and ML analyzed and interpreted the data. HA wrote the initial draft of the paper. SJ, JT and ML revised the paper. All authors critically reviewed and approved the final manuscript.

## Ethics Approval and Consent to Participate

### Consent for Publication

### Competing Interests

“The authors declare no conflict of interest.”

## Additional file legends

**Additional file1: Table S1**. CSSL information showing *Og_MG12* introgression(s) for each line and their localization(s) (chromosome and left-right SSR markers). Target fragments were those selected so as to constitute the whole *Og_MG12* genome with the 60 CSSLs. **Table S2**. Quantification of panicle traits in the CSSLs and parents *Os*_*Caiapó* and *O. glaberrima*_*MG12*. Abbreviations: RL, Rachis Length; PBN, Primary Branch Number; PBL, Primary Branch Length; SBN, Secondary Branch Number; SpN, Spikelet Number; SBN/PBN, ratio of SBN and PBN. **Table S3**. Means and broad sense heritability for each scored trait. **Table S4**. Variation and descriptive statistics of the scored traits in the CSSL population and the parents. **Table S5**. Significant line*trait associations observed in the population studied. **Table S6**. QTL and GWAS sites colocalized with the QTLs detected in this study. **Table S7**. Metrics of genome quality obtained for *Os_Caiapó* and *Og_MG12* genome ONT sequencing assemblies. **Table S8**. Summary of the positions (in bp) of structural variations observed in the QTLs detected. **Table S9**. Gene synteny and description of loci annotated between the RM10 and RM420 markers in *Os_Caiapó* (RAP_db and MSU annotations) and *Og_MG12* (RAP_db, MSU and OglaRS2). Genes highlighted in yellow are absent in the q_7 region of *Og_MG12*, genes highlighted in green are annotated in the *Og_MG12* q_7 region but are absent in the *Os_Caiapó* genome, genes highlighted in light green are duplicated in the *Og_MG12* q_7 region and genes highlighted in pink are annotated in the *Og_MG12* q_7 region, are absent in the *Os_ Caiapó* q_7 region but are present in other chromosomes in the *Os_Caiapó* genome. **Table S10**. Amino acid modifications in proteins encoded by candidate genes in q_7. **Table S11**. TFBS variations in the promoters of candidate genes present in q_7. Abbreviations: EIN3, ETHYLENE-INSENSITIVE-LIKE3; C2H2, C2H2-Type Zinc finger; AP2, AP2-ERF; bHLH, basic helix-loop-helix; TCP, TEOSINTE-BRANCHED1; SBN, SQUAMOSA BINDING PROTEIN; NY-YB, nuclear factor Y; HB, HOMEODOMAIN; MADS, MADS-BOX; bZIP, basic leucine zipper; TBP, TATA-box-binding.

**Additional file2: Figure S1**. CSSL Finder screenshots showing graphical representations of the genotypes of the 60 BC_3_DH lines along with corresponding data for each evaluated panicle trait. Abbreviations: A. RL, Rachis Length; B. SBN, Secondary Branch Number; C. SpN. Spikelet Number; D. PBN, Primary Branch Number; E. PBL, Primary Branch Length; F. TBN, Tertiary Branch Number. The 12 chromosomes are displayed vertically. They are covered by 200 evenly dispersed SSR markers. The genotypes of the individual lines are displayed horizontally. Shading indicates the allelic composition of chromosomes. Light gray areas represent the *Os_Caiapó* genetic background, black areas represent the *Og_MG12* chromosome segments, dark gray areas represent the heterozygous segments and blue areas correspond to missing data. On the right, solid color bars indicate the values of the panicle traits tested for each line. At the bottom of each graph, the dotted line indicates the statistical threshold of the F-test for the evaluated panicle trait.

**Additional file3: Figure S2**. Genetic and phenotypic description of CSSLs showing the extreme significant differences for the panicle trait analyzed (A) SBN/PBN, (B) SpN, (C) PBN, (D) PBL and (E) TBN. Upper subpanel: Chromosome graphical representation of *Og_MG12* introgression positions in the CSSL. Position of the SSR markers is indicated in Mbp. Lower subpanel: boxplots of the phenotypic variation observed in Year 1 and Year 2 in these lines. Each point represents the phenotypic value for one panicle. Statistical significance (*t*-test *p*-values) between *Os*_*Caiapó* and each line for the panicle morphological trait is indicated as follows: NS if the test is non-significant; *p-values < 0.05; **<0.01; ***<0.001. Abbreviations: PBN, primary branch number; PBL, primary branch length; TBN, tertiary branch number; SBN/PBN, ratio between secondary branch and primary branch numbers; SpN, Spikelet number.

**Additional file4: Figure S3**. Phenotypic description of BC_4_ lines to dissect the effects of QTLs on panicle traits. Boxplot of the phenotypic variation observed in *Os_Caiapó* (yellow), *Og_MG12* (green), and BC_4_ (blue) lines in greenhouse (2021). Each point represents the phenotypic value for one panicle. Statistical significance (*t*-test *p*-values) between *Os_Caiapó* and each line for the panicle morphological traits is indicated as follows: ** *p*-values <0.01; ***<0.001. Abbreviations: PBN, primary branch number; PBL, primary branch length; TBN, tertiary branch number; SBN/PBN, ratio between secondary branch and primary branch numbers; SpN, spikelet number.

**Additional file5: Figure S4**. Dotplots of the minimap2 alignment (implemented in D-GENIES web facilities) of panicle trait-related QTLs detected between the *Os_Caiapó* and *Og_MG12* genomes and between each of the latter aligned against *O. sativa* cv. *Nipponbare* as a reference genome. QTL coordinates in each genome are indicated on axis X and Y. Dot colors are relative to the identity value (I) which is a BLAST-like alignment identity (I=Number of bases, including gaps per number of matching bases in the bases).

**Additional file6: Figure S5**. Physical map positions of detected QTLs and colocalizations with known genes.

## Notes

### Competing Interest Statement

The authors have declared no competing interest.

