## Supplemental Figures for "African rice (*Oryza glaberrima*) genomic introgressions impacting upon panicle architecture in Asian rice (*O. sativa*) lead to the identification of key QTLs": AdditionalFile2-6 Figures.pdf

### Additional file 2: Figure S1.

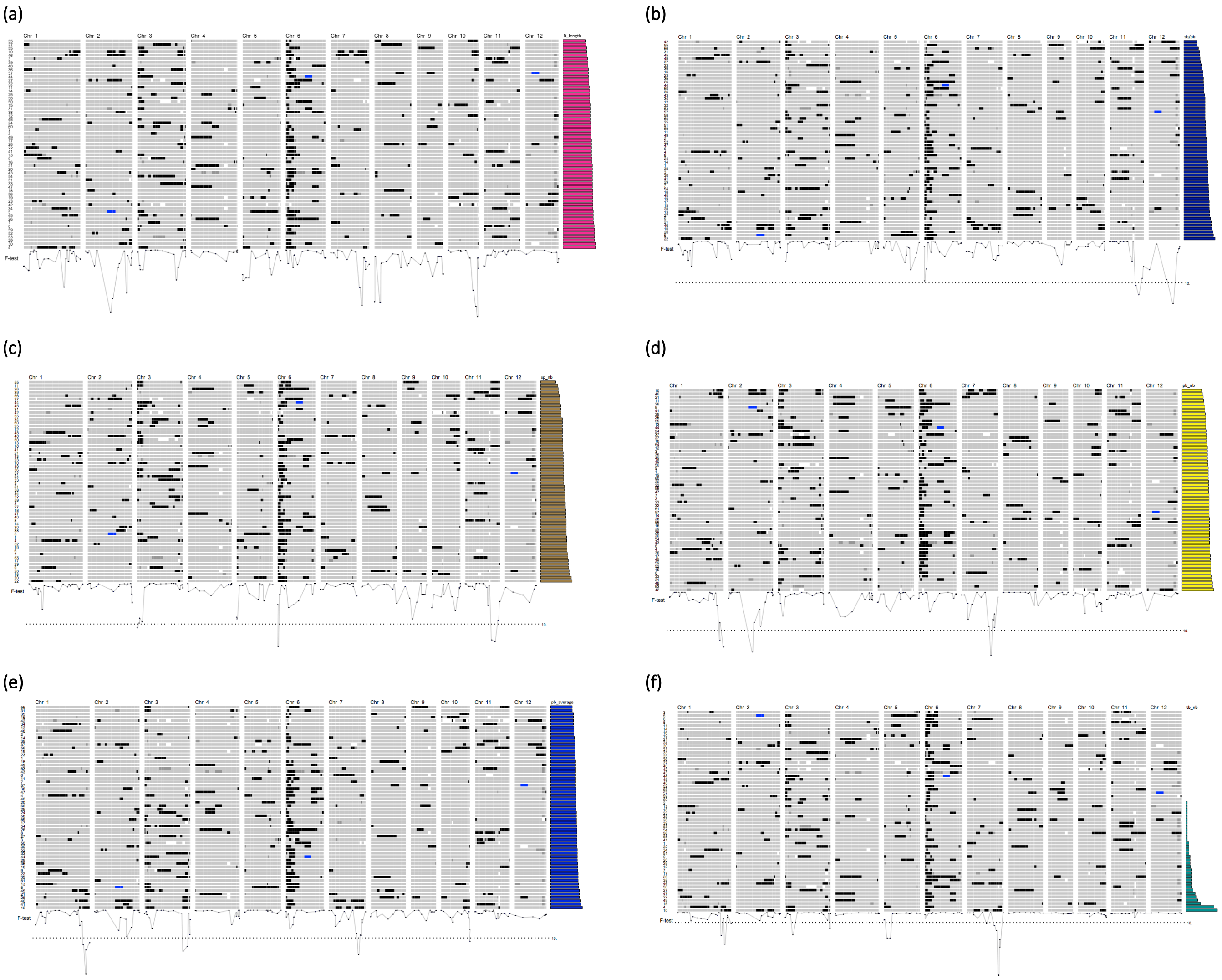

**Additional file 2: Figure S1.** CSSL Finder screenshots showing a graphical representations of the genotypes of the 60 $BC_3DH$ lines along with corresponding data for each evaluated panicle trait. **(a)** RL, Rachis Length; **(b)** SBN, Secondary Branch Number; **(c)** SpN. Spikelet Number; **(d)** PBN, Primary Branch Number; **(e)** PBL, Primary Branch Length; **(f)** TBN, Tertiary Branch Number. The 12 chromosomes are displayed vertically. They are covered by 200 evenly dispersed SSR markers. The genotypes of the individual lines are displayed horizontally. Shading indicates the allelic composition of chromosomes. Light gray areas represent the *Os_Caiapó* genetic background, black areas represent the *Og_MG12* chromosome segments, dark gray areas represent the heterozygous segments and blue areas correspond to missing data. On the right, solid color bars indicate the values of the panicle traits tested for each line. At the bottom of each graph, the dotted line indicates the statistical threshold of the F-test for the evaluated panicle trait.

**Additional file 3: Figure S2.**

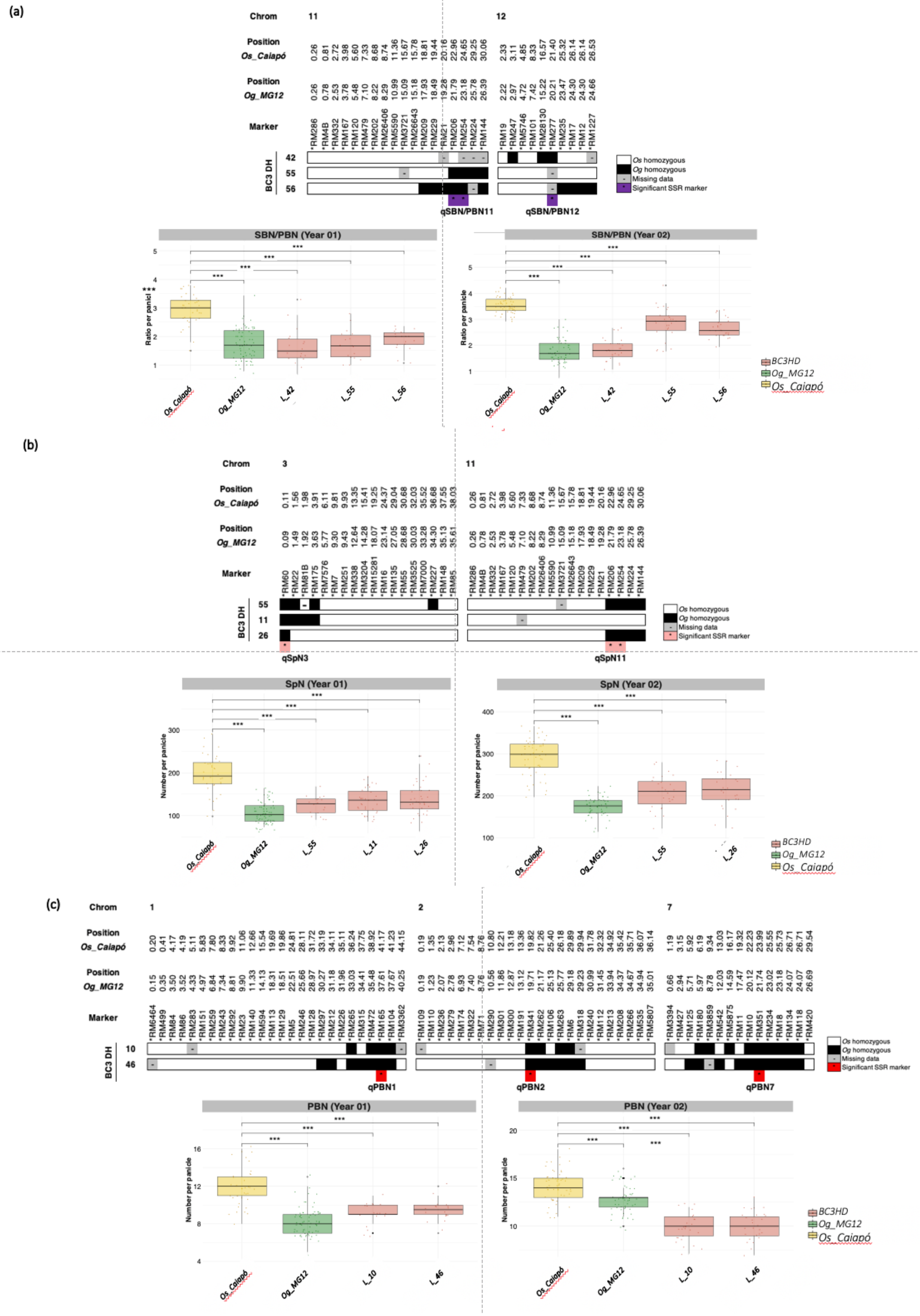

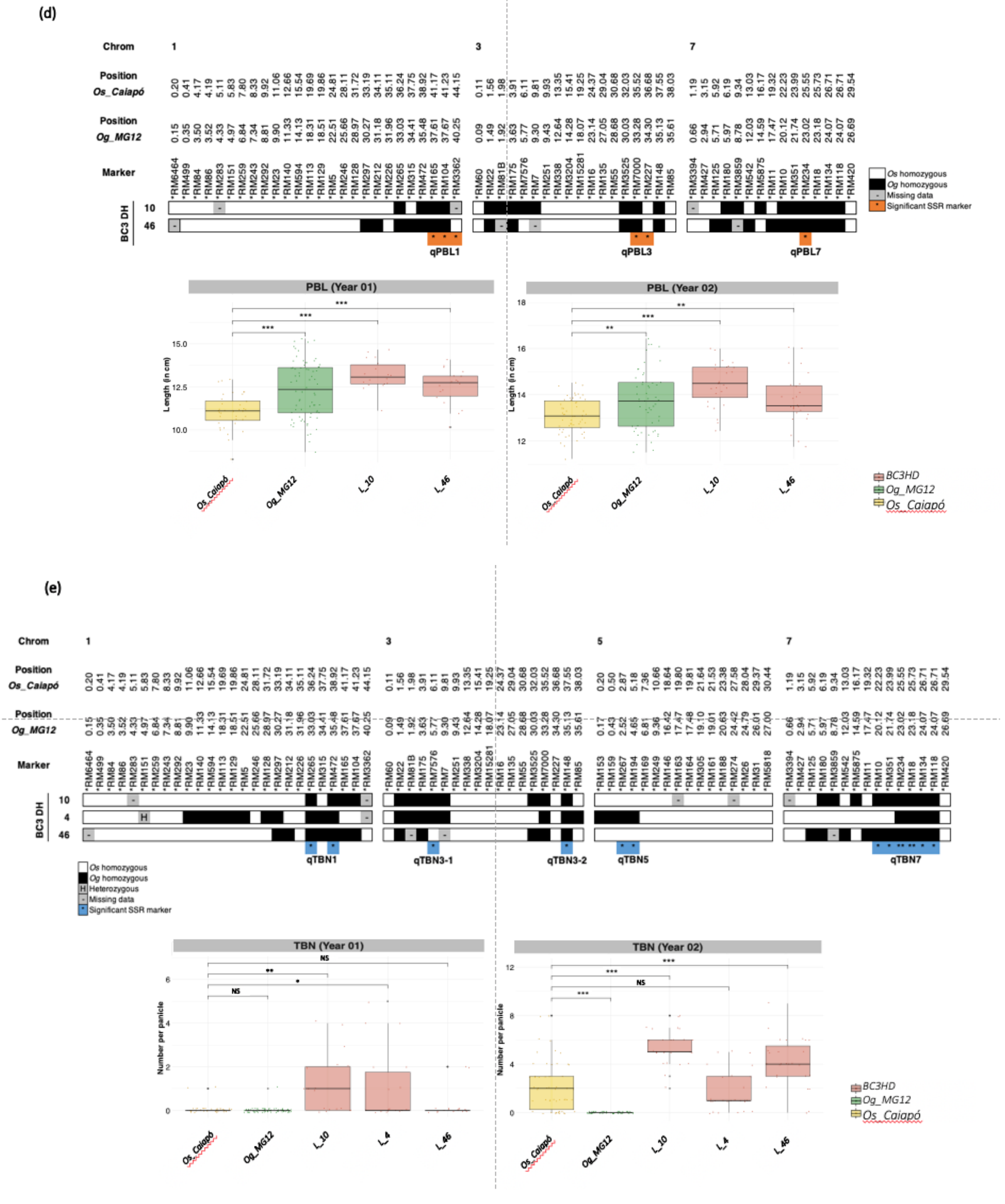


**Additional file 3: Figure S2.** Genetic and phenotypic description of CSSLs showing the extreme significant differences for the panicle trait analyzed **(a)** SBN/PBN; **(b)** SpN; **(c)** PBN; **(d)** PBL; and **(e)** TBN. Upper subpanel: Chromosome graphical representation of *Og_MG12* introgression positions in the CSSL. Position of the SSR markers is indicated in Mbp. Lower subpanel: boxplots of the phenotypic variation observed in Year 1 and Year 2 in these lines. Each point represents the phenotypic value for one panicle. Statistical significance (*t*-test *p*-values) between *Os_Caiapó* and each line for the panicle morphological trait is indicated as follows: NS if the test is non-significant; *p-values < 0.05; **<0.01; ***<0.001. Abbreviations: PBN, primary branch number; PBL, primary branch length; TBN, tertiary branch number; SBN/PBN, ratio between secondary branch and primary branch numbers; SpN, Spikelet number.

### Additional file 4: Figure S3.

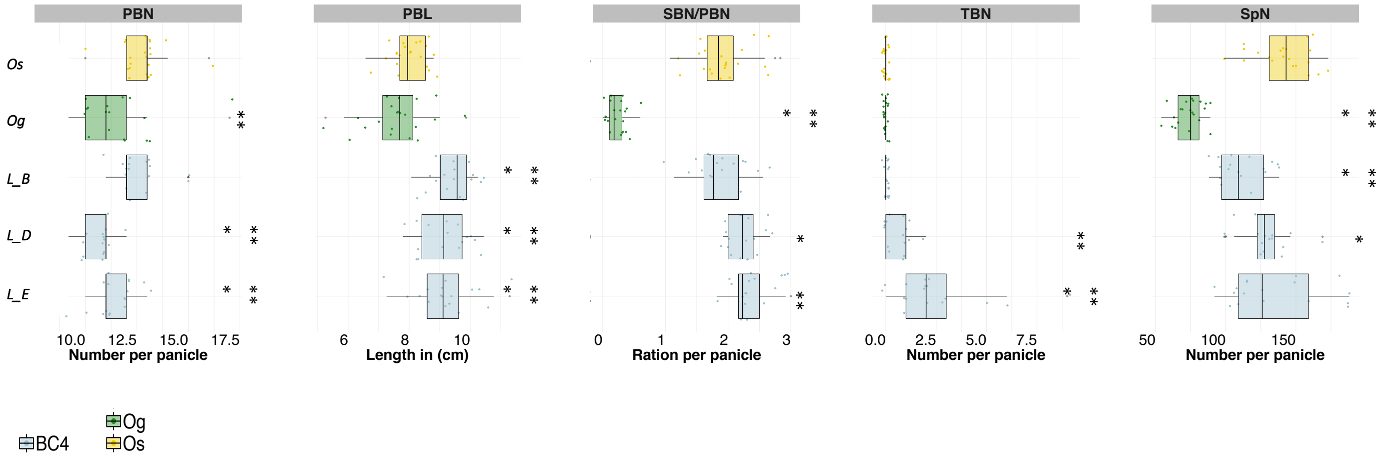


**Additional file 4: Figure S3.** Phenotypic description of $BC_4$ lines to dissect the effects of QTLs on panicle traits. Boxplot of the phenotypic variation observed in *Os_Caiapó* (yellow), *Og_MG12* (green), and $BC_4$ (blue) lines in greenhouse (2021). Each point represents the phenotypic value for one panicle. Statistical significance (*t*-test *p*-values) between *Os_Caiapó* and each line for the panicle morphological traits is indicated as follows: ** *p*-values <0.01; ***<0.001. Abbreviations: PBN, primary branch number; PBL, primary branch length; TBN, tertiary branch number; SBN/PBN, ratio between secondary branch and primary branch numbers; SpN, spikelet number.

**Additional file 5: Figure S4.**

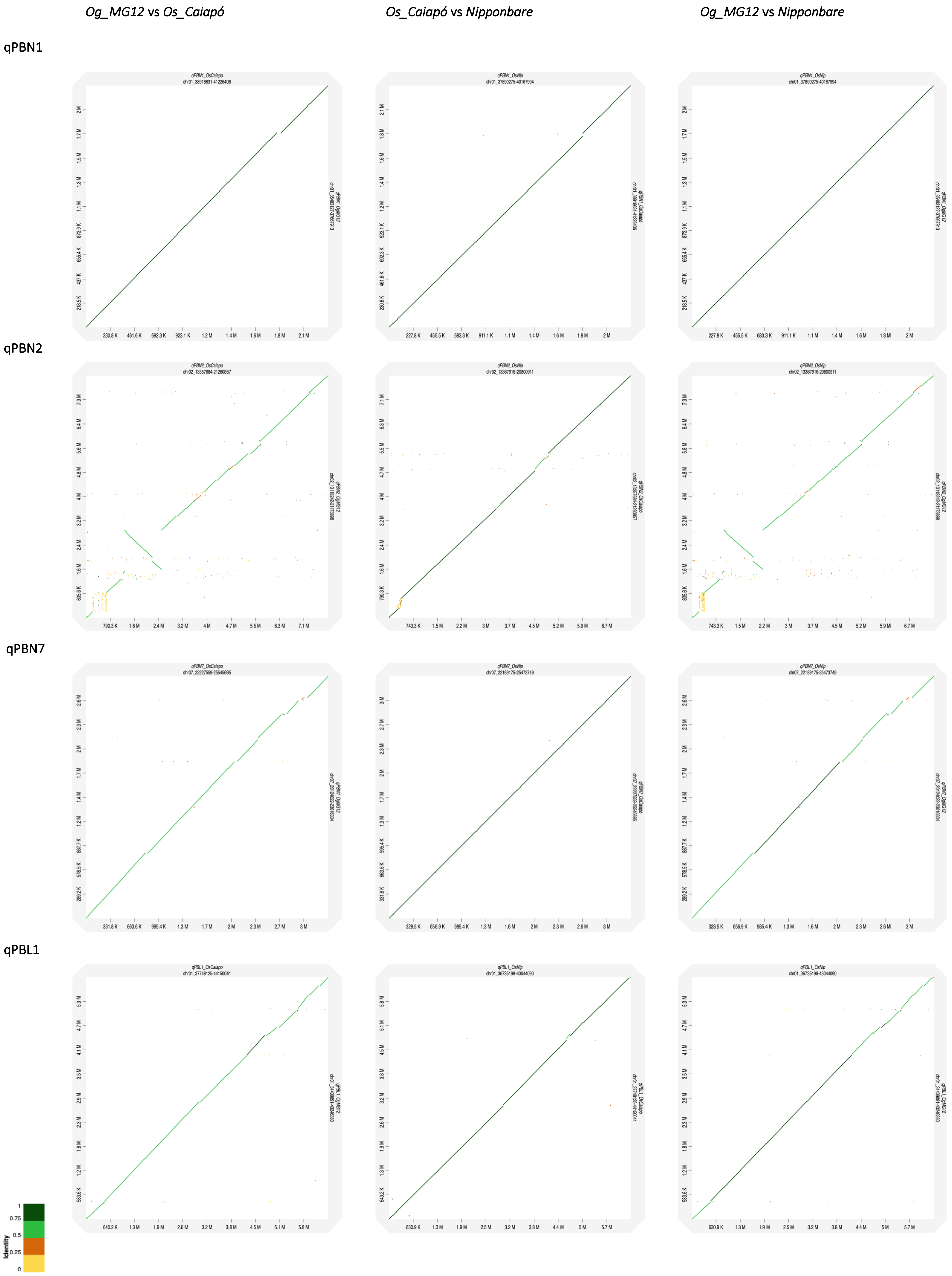

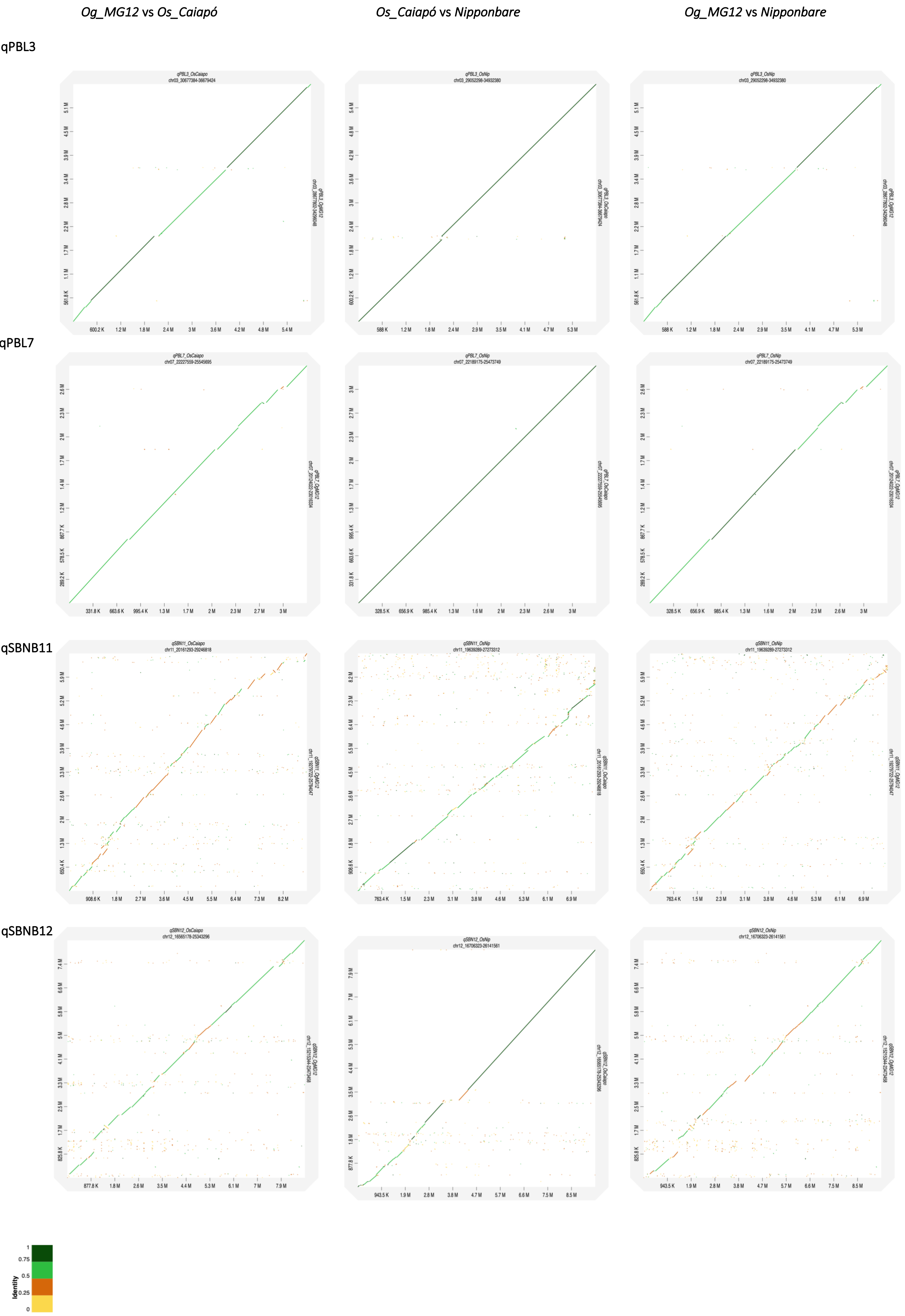

Og_MG12 vs Os_Caiapó
Os_Caiapó vs Nipponbare
Og_MG12 vs Nipponbare
qPBL3
qPBL7
qSBNB11
qSBNB12
Identity
1
0.75
0.5
0.25
0

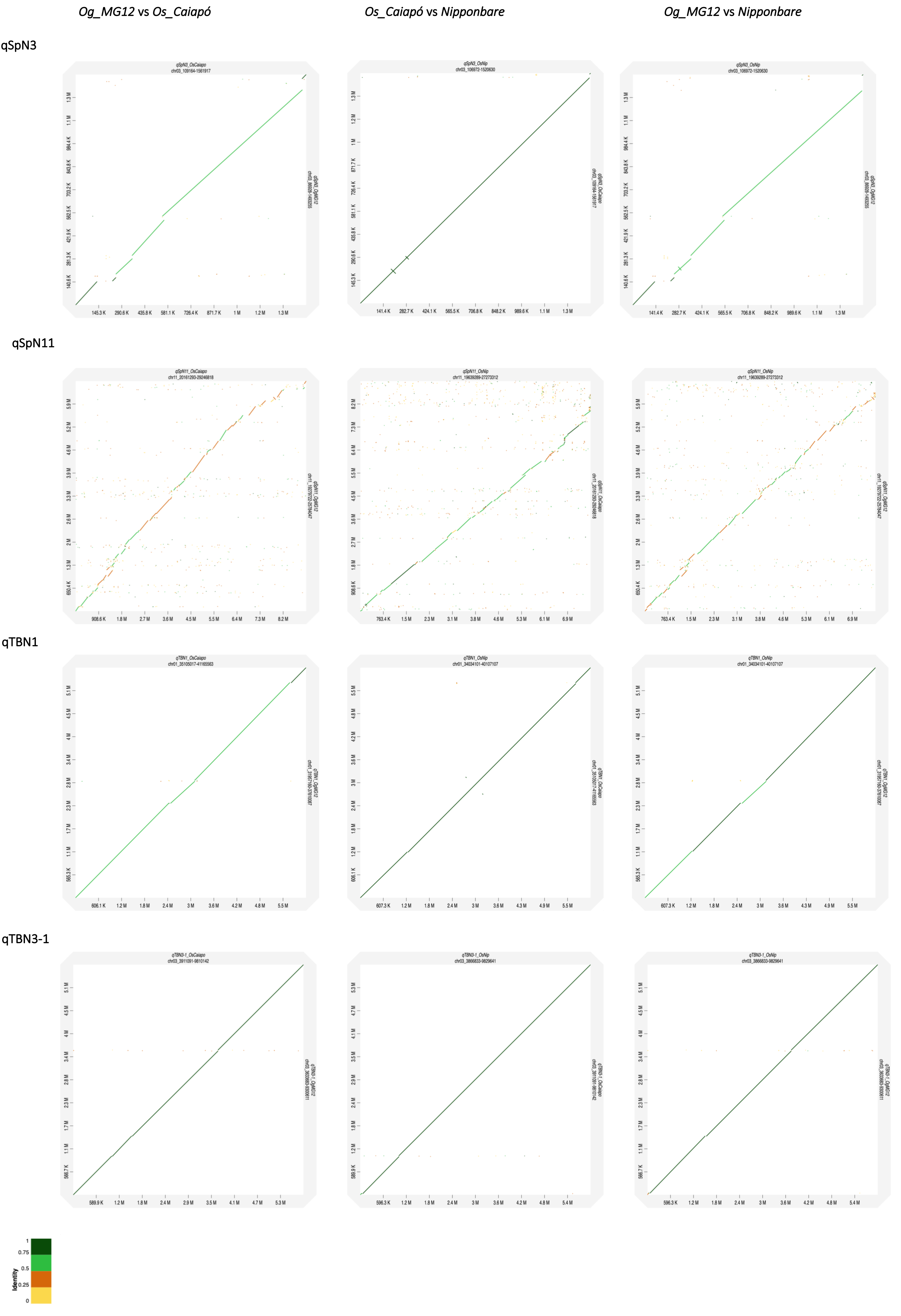

Og_MG12 vs Os_Caiapó
Os_Caiapó vs Nipponbare
Og_MG12 vs Nipponbare
qSpN3
qSpN11
qTBN1
qTBN3-1
Identity
1
0.75
0.5
0.25
0

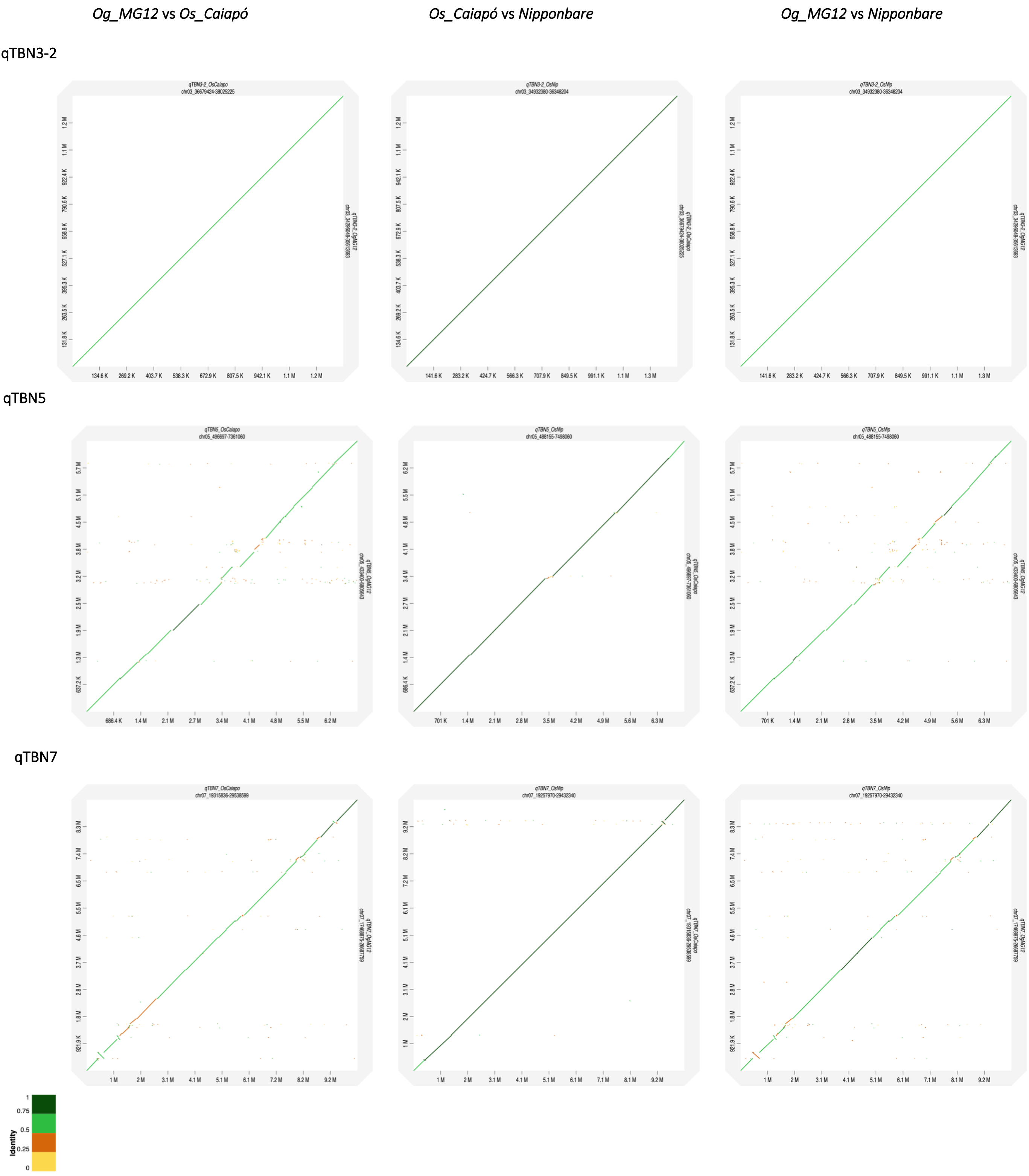


**Additional file 5: Figure S4.** Dotplots of the minimap2 alignment (implemented in D-GENIES web facilities) of panicle trait-related QTLs detected between the *Os_Caiapó* and *Og_MG12* genomes and between each of the latter aligned against *O. sativa* cv. *Nipponbare* as a reference genome. QTL coordinates in each genome are indicated on axis X and Y. Dot colors are relative to the identity value (I) which is a BLAST-like alignment identity (I=Number of bases, including gaps per number of matching bases in the bases). Correction to make in the figure (top right): Nipponabre -> Nipponbare

**Additional file 6: Figure S5.**

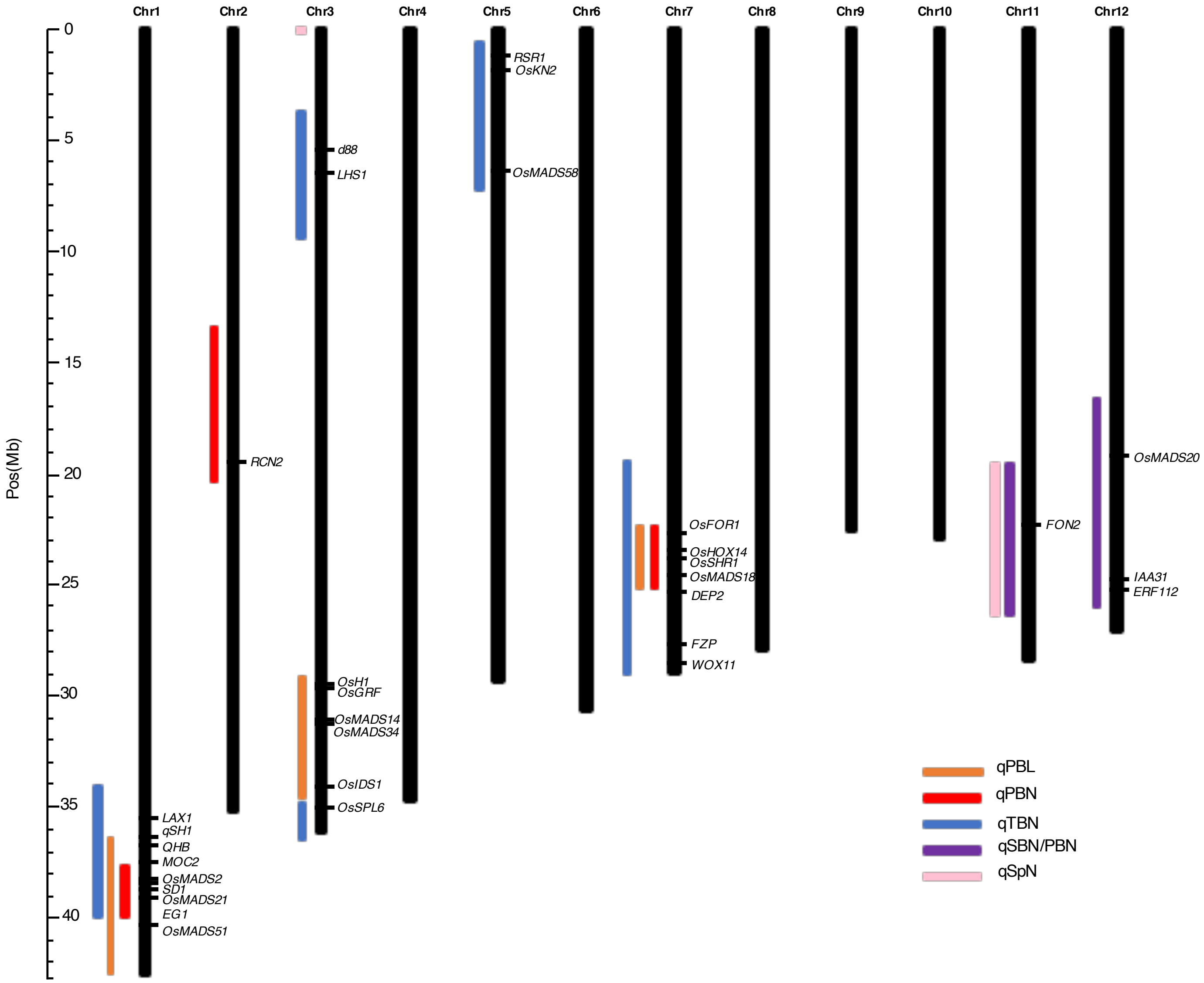


**Additional file 6: Figure S5** : Physical map positions of detected QTLs and colocalizations with known genes.
